# Dynamics of dN/dS within recombining bacterial populations

**DOI:** 10.1101/2025.09.09.675256

**Authors:** Zhiru Liu, Benjamin H. Good

## Abstract

The ratio of nonsynonymous to synonymous substitutions (dN/dS) encodes important information about the selection pressures acting on protein-coding genes. In bacterial populations, dN/dS often declines with the sequence divergence between strains, but the mechanisms responsible for this broad empirical trend are still debated. Existing models have primarily focused on de novo mutations, overlooking the older genetic variants that are continually introduced through horizontal gene transfer and recombination. Here we introduce a phenomenological model of dN/dS in recombining populations of bacteria, which allows us to disentangle the effects of recombination among pairs of closely related strains. We find that clonally inherited regions of the genome exhibit consistently higher dN/dS ratios, and that the accumulation of recombined segments can quantitatively explain the majority of the decline in dN/dS. We use these observations to re-examine models of purifying selection and adaptive reversion in human gut bacteria, and uncover evidence for widespread weak selection at a large fraction of protein coding sites. Our findings show that horizontal gene transfer can be an important factor in shaping genome-wide patterns of selective constraint, and raise new questions about the effectiveness of natural selection in complex bacterial populations.

## Introduction

Patterns of DNA sequence conservation play a central role in modern evolutionary biology. In protein-coding genes, the degree of conservation is often quantified by comparing the rates of non- synonymous and synonymous substitutions (dN/dS) (1). Under the assumption that synonymous mutations are effectively neutral, the dN/dS ratio provides a simple readout of the net direction of natural selection (2, 3): values less than one imply that many nonsynonymous variants have been eliminated by negative selection, while ratios greater than one imply that some of the observed substitutions were positively selected. This simple principle has widespread applications throughout evolutionary biology, from the identification of functionally important DNA sequences (4–6) to mapping the causative loci responsible for driving short-term evolutionary adaptation (7–10).

While originally developed for comparisons between species, dN/dS is also routinely applied to genomes from the same species (11, 12). In this case, the genetic differences between these conspecific genomes represent polymorphisms that are circulating within the larger species population. An extensive literature has shown that the ratios of non-synonymous and synonymous variants at different allele frequencies can enable quantitative inferences of the underlying selection coefficients acting on new mutations (12–16), by partitioning the action of natural selection over different historical timescales. A similar approach is common in bacterial populations, where dN/dS can be computed for pairs of strains with different degrees of genetic relatedness. A widespread observation within many bacterial species is that dN/dS gradually decreases with increasing genetic divergence between strains, ranging from nearly neutral levels (dN/dS ≈ 1) among the most closely related strains to much lower levels (dN/dS≈0.1) between typical pairs of conspecific isolates (8, 17–29). This robust trend suggests that dN/dS is time-dependent, with selection acting to reduce nonsynonymous variation as divergence time increases.

However, the mechanisms responsible for this time-dependent decay remain under debate. The classical explanation attributes the decline in dN/dS to the finite strength of purifying selection (22, 25, 28): weaker deleterious mutations take some time to be purged from the population, leading to an enrichment of non-synonymous variants among the most recently diverged strains. A more recent alternative proposes that the higher dN/dS values at short times arise from locally adaptive substitutions that undergo reversions on longer timescales, producing a long-term trend that is reminiscent of classical evolutionary constraint (30). While these two models rely on fundamentally different mechanisms, previous work suggests that they can both reproduce the observed dN/dS decay trends in multiple species of human gut bacteria (28–30), making them difficult to distinguish on the basis of these aggregate statistics alone.

Crucially, both of these existing explanations overlook a key process of bacterial evolution, where homologous recombination replaces a portion of the genome with a corresponding segment from another donor strain (31). Previous studies have shown that recombined regions in recently diverged genomes often exhibit lower dN/dS than clonal regions (24). This suggests that recombination could mimic the effects of natural selection by introducing older genetic variants that have already been filtered by selection. Given the widespread prevalence of homologous recombination within many bacterial species (32–35), this raises the possibility that the temporal trend in dN/dS could be strongly shaped by the accumulation of recombination events over time.

Here, we investigate this hypothesis by studying the impact of bacterial recombination on the dynamics of dN/dS. We show that a simple model in which recombination introduces divergent segments from unrelated donors can quantitatively recapitulate the observed decline in genomewide dN/dS values within a large panel of human gut bacteria. After correcting for the confounding effects of recombination, we show how the residual dN/dS dynamics in clonally inherited regions can constrain existing models of purifying selection and adaptive reversion. These findings highlight the dominant role of recombination in shaping genome-wide patterns of selective constraint, and offer new insights into the dynamics of selection in natural bacterial populations.

## Results

### Quantifying the impact of homologous recombination on dN/dS dynamics

To illustrate how horizontal gene transfer and recombination can alter the temporal dynamics of dN/dS, we consider a simple conceptual model based on the genealogies of the underlying strains (33, 34, 36, 37; Fig. 1). We imagine a genome-wide comparison between two conspecific strains that descended from a single cell division event exactly *T* generations in the past (Fig. 1A). The genetic differences between these bacterial genomes will arise from two primary sources: (i) de novo mutations that are inherited through clonal descent, and (ii) pre-existing variants that are introduced via homologous recombination with other strains (Fig. 1B).

**Figure 1:**
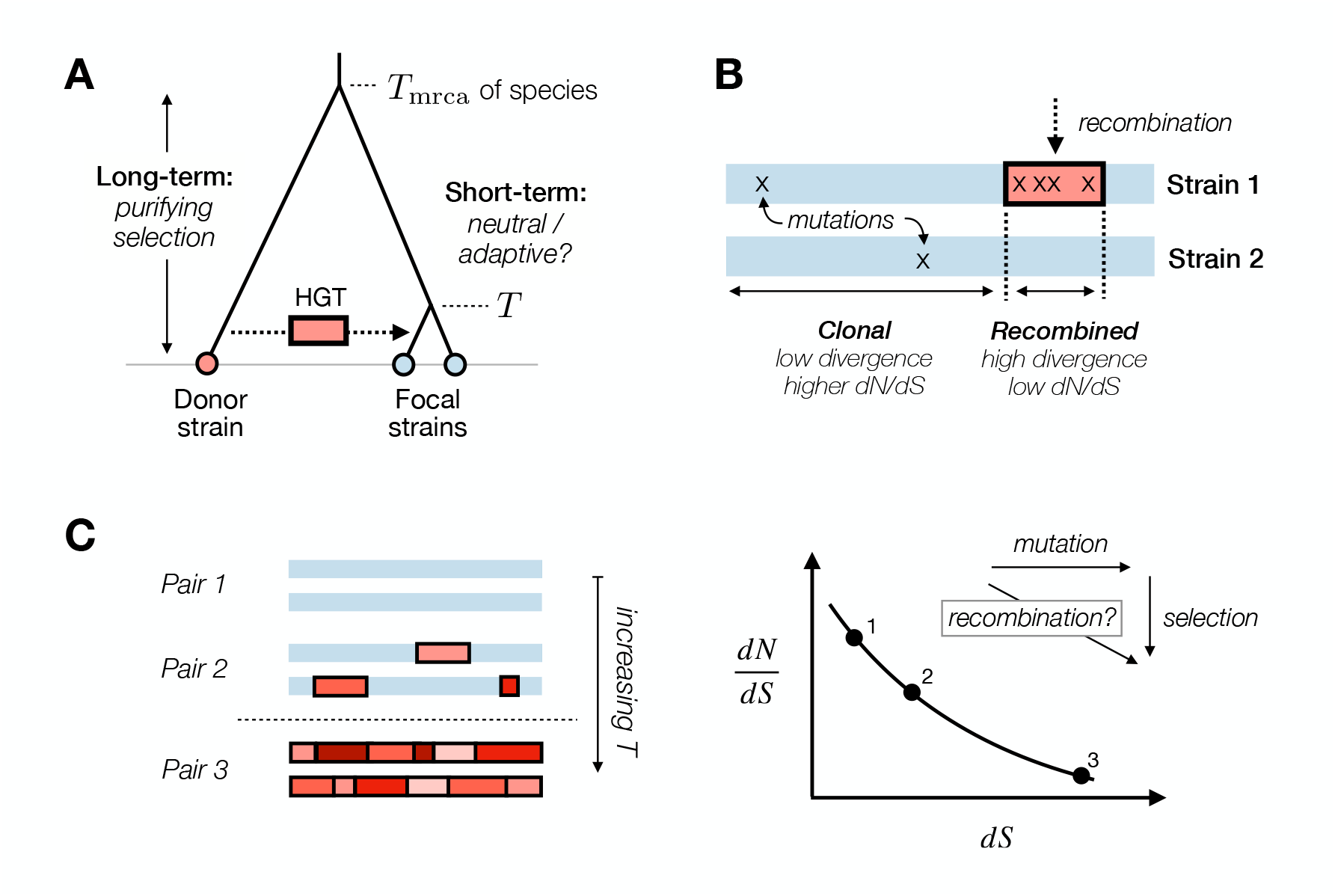
Modeling the impact of homologous recombination on dN/dS between conspecific strains of bacteria. (A) Pairs of recently diverged strains that shared a common ancestor *T* generations ago can acquire homologous DNA segments from more distantly related strains of the same species, which shared a common ancestor ∼*T*_mrca_ generations ago. (B) Recombined segments (red) introduce a high density of older genetic variants that have already been filtered by natural selection (low dN/dS), while clonally inherited regions (blue) exhibit lower genetic divergence and higher dN/dS from more recent de novo mutations. (C) The observed values of dN and dS in different pairs of strains (left) are influenced by their total proportion of clonal (blue) versus recombined (red) regions. The steady accumulation of recombination events produces a downward trend in dN/dS as strains diverge, potentially obscuring the underlying dynamics of natural selection.

Under the assumption that synonymous mutations are effectively neutral, de novo mutations will generate single nucleotide differences (SNVs) at a characteristic rate of *d*_*S*_ ≈ 2*µT*, where *µ* is the corresponding mutation rate in units of mutations per site per generation. In contrast, the synonymous divergence introduced by recombination will depend on the net rate of recombination per site (*R*), as well as the genetic divergence of the imported segments. Previous work has shown that these recombined DNA fragments often originate from unrelated donor strains (33, 34, 36, 37), which carry an average synonymous divergence of 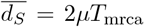 (Fig. 1A). When *T* ≪ 1*/R* ≪ *T*_mrca_, the introduction of SNVs through recombination can substantially inflate *d*_*S*_ relative to the baseline expectation from de novo mutations alone. This hybrid divergence process continues until the majority of the clonal backbone has been overwritten by homologous recombination (*T* ≳ 1*/R*), after which *d*_*S*_ rapidly converges to the steady-state value 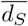.

This phenomenological model can also be extended to incorporate non-synonymous SNVs. In clonal regions of the genome, non-synonymous divergence accumulates at a time-dependent rate *d*_*N*_ (*T*), whose detailed shape will depend on the underlying model of selection (described in more detail in the following section). Recombined segments, by contrast, will carry an average nonsynonymous divergence of 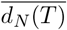 which can be estimated from the standing genetic diversity between unrelated pairs of strains (i.e. those with *T* ≳ 1*/R*). When *d*_*N*_ (*T*) is a non-linear function of time, the value of dN/dS in the clonal regions of the genome will generally differ from the recombined segments. This reflects the fact that mutations on recombined segments have already been circulating in the population for a longer period of time, providing more opportunities for natural selection to act (Fig. 1A,B). As genomes accumulate more recombination events, the growing contribution from the recombined segments can alter the genome-wide average of dN/dS, similar to how they inflate the overall genetic divergence (Fig. 1C). Notably, this recombination process can generate a smoothly declining trend in the genome-wide dN/dS over time, even if the dN/dS ratios are largely constant within both the clonal and recombined regions.

To test whether this hypothesis could explain the dN/dS dynamics in empirical data, we leveraged a previously curated dataset of commensal human gut bacteria (28), containing 5,416 reconstructed strain genotypes from 43 prevalent species (SI Section 1). The genetic distances between these conspecific strains span nearly three orders of magnitude, from a typical value of *d*_*S*_ ∼ 1% to just a handful of detected SNVs (*d*_*S*_ ≲ 10^−5^); the corresponding *d*_*N*_ */d*_*S*_ levels vary over an order of magnitude as well (Fig. 2B). In a recent study (34), we used a hidden Markov model (HMM) approach to label individual recombination events and clonally inherited regions within the most closely related genomes in this dataset (Fig. 2A). Since our HMM method relies on the spatial distribution of synonymous variants, these labels should be independent of the corresponding *d*_*N*_ levels. The conceptual model in Fig. 1 then leads to a clear prediction for two different sources of data: if recombination is the primary driver of the dN/dS decay in Fig. 2B, then stratifying sites by their recombination status should reveal distinct dN/dS trajectories.

**Figure 2:**
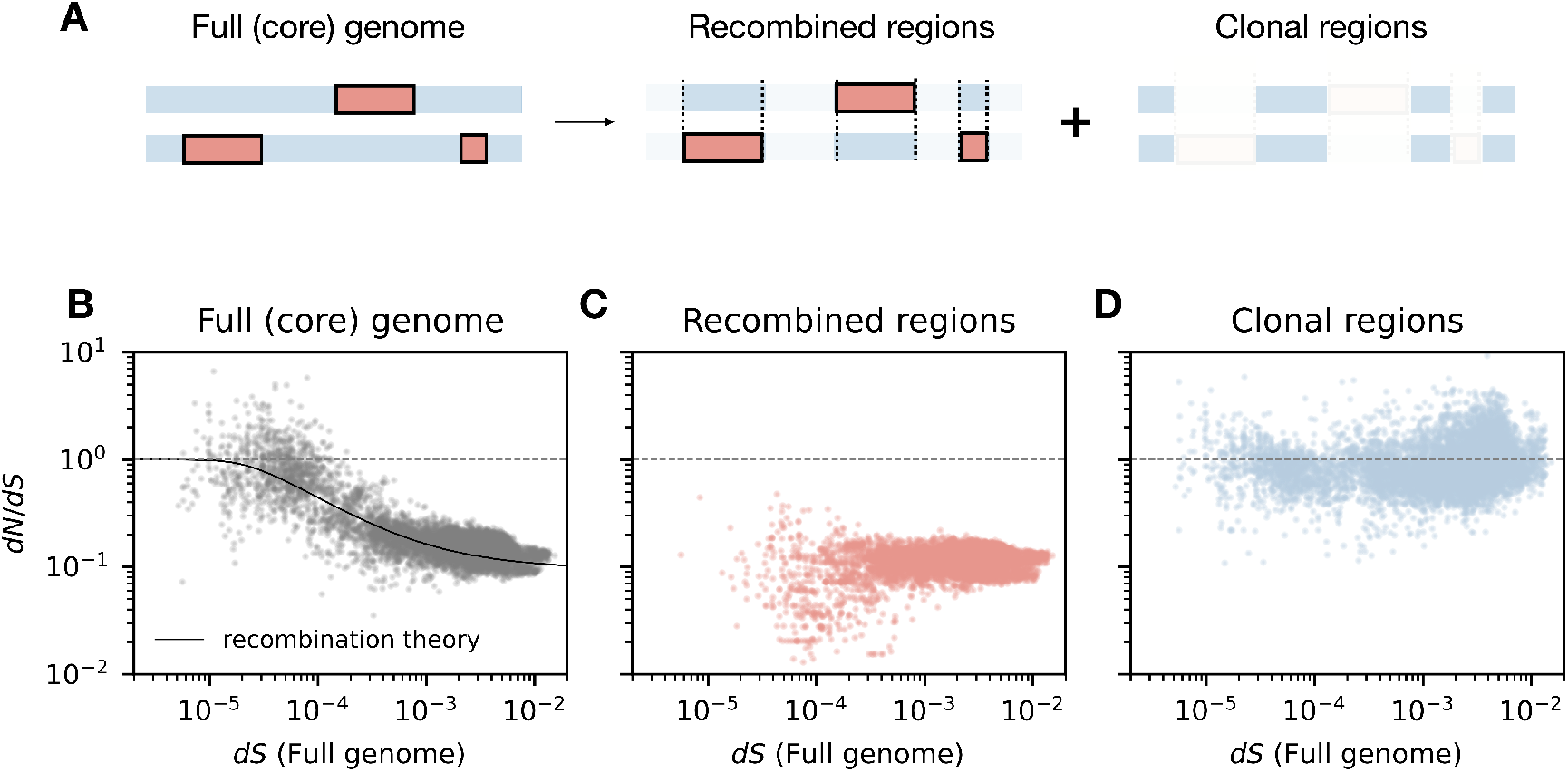
Temporal dynamics of dN/dS reflect the accumulation of homologous recombination events in human gut commensals. (A) Schematic illustration of how recombination events are isolated. Recombined regions (red blocks) and clonal regions (blue blocks) are identified based on the local density of synonymous SNVs. dN/dS values are then computed separately for each region type, as shown in panels C and D. (B) Genome-wide dN/dS dynamics in 29 human gut bacterial species reproduced from Ref. 28. Each point represents the genome-wide dN/dS value estimated from non-degenerate (1D) and fourfold-degenerate (4D) sites for an individual pair of closely related strains (SI Section 2). The solid line shows the prediction of the recombination model in SI Section 3, while the dashed line indicates the neutral expectation (dN/dS=1). (C-D) Analogous versions of panel B showing dN/dS values computed within the recombined (C) or clonal (D) of the genome separately.

Consistent with this prediction, we found that the dN/dS dynamics in our dataset dramatically differ between clonal and recombined segments (Fig. 2C,D). Within recombined regions, dN/dS is consistently low (*d*_*N*_ */d*_*S*_ ~ 0.1), closely matching the values observed between unrelated pairs of strains. In contrast, dN/dS remains close to one within the clonal regions, even as genome-wide divergence increases by several orders of magnitude.

To demonstrate how these local patterns give rise to the full genome-wide trend in Fig. 2B, we formulated a simple mathematical model of dN/dS accumulation that accounts for the growing contribution of recombined segments over time (Fig. S1; SI Section 3). Our phenomenological model simply averages the observed dN/dS values in the clonal and recombined regions, and weights them by the empirically determined fraction of the genome covered by recombination at each divergence level dS. As a result, the model has no additional free parameters that require fitting to the observed trend.

Despite its simplicity, we found that this mixture model quantitatively reproduces the observed dN/dS decay across the full range of divergence levels in Fig. 2B. In particular, it captures the characteristic transition from high to low dN/dS values at intermediate levels of synonymous divergence. In previous dN/dS models, the location of this transition was related to the underlying timescales of the natural selection (e.g., the selection coefficient in the purifying selection model of Ref. 28, or the reversion rate in the adaptive reversion model in Ref. 30). In our present model, this transition naturally emerges from the increasing genomic footprint of recombination, and occurs around the point when recombined segments begin to dominate the genome (Fig. 1C).

To explore whether these results extend beyond human gut bacteria, we also re-analyzed the dN/dS dynamics in 110 strains of *Staphylococcus aureus* (SI Section 7). *S. aureus* is a common bacterial pathogen with a distinct ecology and recombination dynamics compared to many gut commensals (38–42). Although clinically isolated strains of *S. aureus* are largely clonal in their recent evolutionary history, three large recombination events have been previously reported in sequence types ST34, ST239 and ST582 (42–44). We used these previously identified events to stratify the genome into clonal and recombined regions as above, and analyzed the corresponding dN/dS patterns in an analogous manner (Fig. S2). Once again, we found that dN/dS dynamics separate into distinct trends by genomic region, consistent with the predictions of our basic recombination model. These results suggest that recombination may be a general mechanism underlying the frequently observed temporal trend in dN/dS across diverse bacterial species.

### Estimating the strength of purifying selection from recombination-corrected dN/dS

While the mixture model in Fig. 2B can recapitulate the overall shape of the dN/dS curve, it still requires external estimates of the dN/dS levels in each region (Fig. 2C,D). These levels are set by the selective forces that act on nonsynonymous variants as they circulate through the broader population, which ultimately produce the low dN/dS values on a typical recombined segment. To probe these dynamics without the confounding influence of recombination, we can use the labels in Fig. 2 to construct a “corrected” dN/dS trajectory, using the local values of *d*_*N*_ and *d*_*S*_ in each genomic region.

The short-term dynamics of dN/dS can be assessed using the clonally inherited regions that are preserved between closely related genome pairs. Fig. 2D shows that the dN/dS levels on the clonal backbone remain close to one even as the nominal dS values span more than three orders of magnitude (10^−5^ *< d*_*S*_ *<* 10^−2^; Fig. 2D). However, because these genome-wide dS levels are inflated by recombination, they overestimate the true divergence time between closely related strains.

To more accurately capture the temporal dynamics within the clonal regions, it is helpful to re-plot the clonal dN/dS values as a function of their *clonal* synonymous divergence 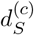 (Fig. 3), which provides a better proxy for the elapsed divergence time between closely related pairs of genomes. The observed clonal divergences span a much narrower range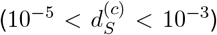, which can help explain the relatively modest decay of dN/dS within these regions. However, the rate of the decay is still about ~10-fold slower than the original data in Fig. 2, highlighting the importance of correcting for recombination. This global trend is also recapitulated at the level of individual species (Fig. S3), with most species showing only a modest decay from *d*_*N*_ */d*_*S*_ ≈ 1 as 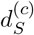 increases. A notable exception is *Alistipes putredinis*, which begins with elevated values (*d*_*N*_ */d*_*S*_ ≈ 2) at the shortest genetic distances and declines below one toward the end of the observable range. Apart from this example in *A. putredinis*, none of the other species showed consistently elevated dN/dS levels within the subset of sites that we are considering.

**Figure 3:**
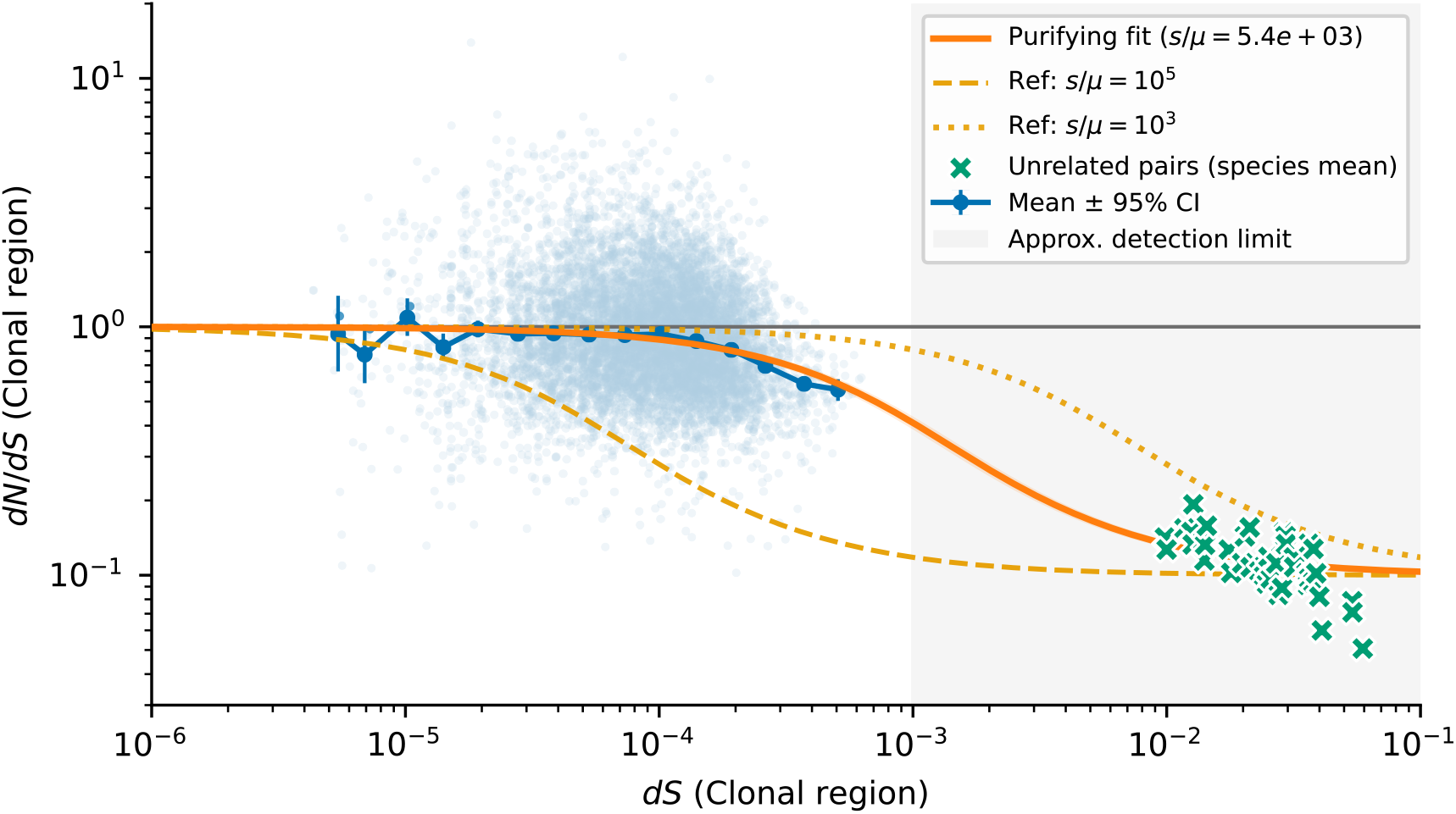
Clonal dN/dS dynamics in human gut bacteria after correcting for recombination. Light blue circles show the clonal *d*_*N*_ */d*_*S*_ values for individual genome pairs (as in Fig. 2D) as a function of their clonal synonymous divergence (SI Section 2). Dark blue circles with error bars indicate the mean and 95% confidence intervals of pooled counts in logarithmically spaced bins of *d*_*S*_ (SI Section 2). Orange solid line shows the best-fit two-class purifying selection model (SI Section 4), with bootstrap confidence intervals narrower than the linewidth. For comparison, the dashed and dotted orange curves show reference predictions with *s/µ* = 10^5^ and 10^3^, respectively, while the horizontal gray line marks the purely neutral expectation (*d*_*N*_ */d*_*S*_ = 1). Green crosses denote the mean *d*_*N*_ */d*_*S*_ and *d*_*S*_ values between fully recombined genome pairs for each individual species, which provide an estimate of the *d*_*N*_ */d*_*S*_ levels when *T* ≈ *T*_mrca_. The shaded vertical band indicates the approximate detection limit (*d*_*S*_ *>* 10^−3^) beyond which most of the genome is overwritten by recombination and clonal tracts cannot be reliably identified (SI Section 1).

For more distantly related strains, recombination events eventually cover the entire genome (Fig. 1C), erasing clonal segments and precluding direct measurement of the clonal dN/dS curve over intermediate timescales. However, we can still estimate the long-term dN/dS levels by examining typical pairs of strains, whose genomes consist entirely of recombined segments with an average synonymous divergence 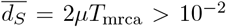. While the spread in this distribution is no longer informative of the differences in the underlying divergence times, the average values still provide a point estimate of the dN/dS levels when *T* ≈ *T*_mrca_. Combining this estimate with the clonal regions above, we arrive at a corrected dN/dS curve with a gap at intermediate times (Fig. 3), corresponding to the transition from clonal to quasi-sexual evolution.

Having corrected for the confounding effects of recombination, we can use the residual dN/dS curves in Fig. 3 to re-examine the support for different models of selection. We begin by considering the standard purifying selection null model, which assumes that nonsynonymous variants are progressively removed over time (22, 28, 30). In this model, the clonal dN/dS curve follows the approximate form,

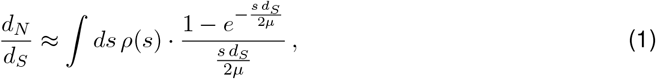

where *ρ*(*s*) is the underlying distribution of fitness costs of nonsynonymous variants (SI Section 4). The key features of the data can be captured by a two-class version of this model, where a fraction of the sites are truly neutral (*s* = 0) and the rest have the same characteristic fitness cost *s*_1_.

Fitting this model to the data in Fig. 3 yields a best-fit value of *s*_1_*/µ* ≈ 10^4^ and a corresponding neutral fraction of ≈10% (SI Section 4). Similar values are obtained when *A. putredinis* is excluded (Fig. S4), indicating that the higher *d*_*N*_ */d*_*S*_ values in this outlier species had little influence on the fitted parameters. These inferred fitness costs are about 10-fold smaller than previous estimates (28–30) obtained from the uncorrected data in Fig. 2 For typical bacterial mutation rates (*µ* ~ 10^−10^ − 10^−9^ per site per generation), the associated selection coefficients are remarkably weak (*s*_1_ ~ 10^−6^ − 10^−5^), and would be difficult to observe on normal experimental timescales. Nevertheless, Fig. 3 shows they are still sufficiently costly that they are eliminated by the time that *T* approaches 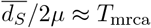.

These weak fitness costs are not an artifact of the two-class model in Fig. 3. Adding a third class of mutations only marginally improves the fit (Fig. S5), and the estimated target size of these mutations is comparatively small (<10%). A more general bound on the strength of selection can be obtained by leveraging the functional form of Eq. 1: the fact that dN/dS remains close to one for clonal divergences as large as 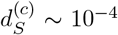 implies that more than 90% of all nonsynonymous mutations must have fitness costs less than *s* ~ 10^5^ · *µ* ≲ 10^−4^ (SI Section 4). At the same time, the fact that dN/dS eventually decays to ≈ 0.1 implies that most of these mutations are eventually purged, which requires that 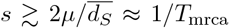. Taken together, these twin constraints imply that the purifying selection model in Eq. 1 can only explain the data if the fitness costs are weak but pervasive, with only a small minority of sites experiencing very strong (*s* ≫ 10^−4^) or nearly neutral (*s* ≪ 1*/T*_mrca_) levels of selective constraint.

### Mutation-type-specific dN/dS dynamics reveal stronger selection against nonsense variants

In the analysis above, all nonsynonymous mutations were aggregated together to compute the temporal dynamics of dN/dS. However, nonsense mutations, which introduce premature stop codons in the gene and often result in a loss of function, are generally expected to be subject to stronger selective pressures than missense mutations. If this is the case, then the model in Eq. 1 predicts that their clonal dN/dS trajectories should diverge from their missense mutation counterparts.

We tested this hypothesis by computing separate dN/dS curves for nonsense and missense mutations, respectively, and examined how each curve varies as a function of the clonal synonymous divergence 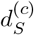 (Fig. 4). The dN/dS decay of missense mutations closely resembled the aggregate results above, as expected given that they constitute the vast majority of nonsynonymous sites.

**Figure 4:**
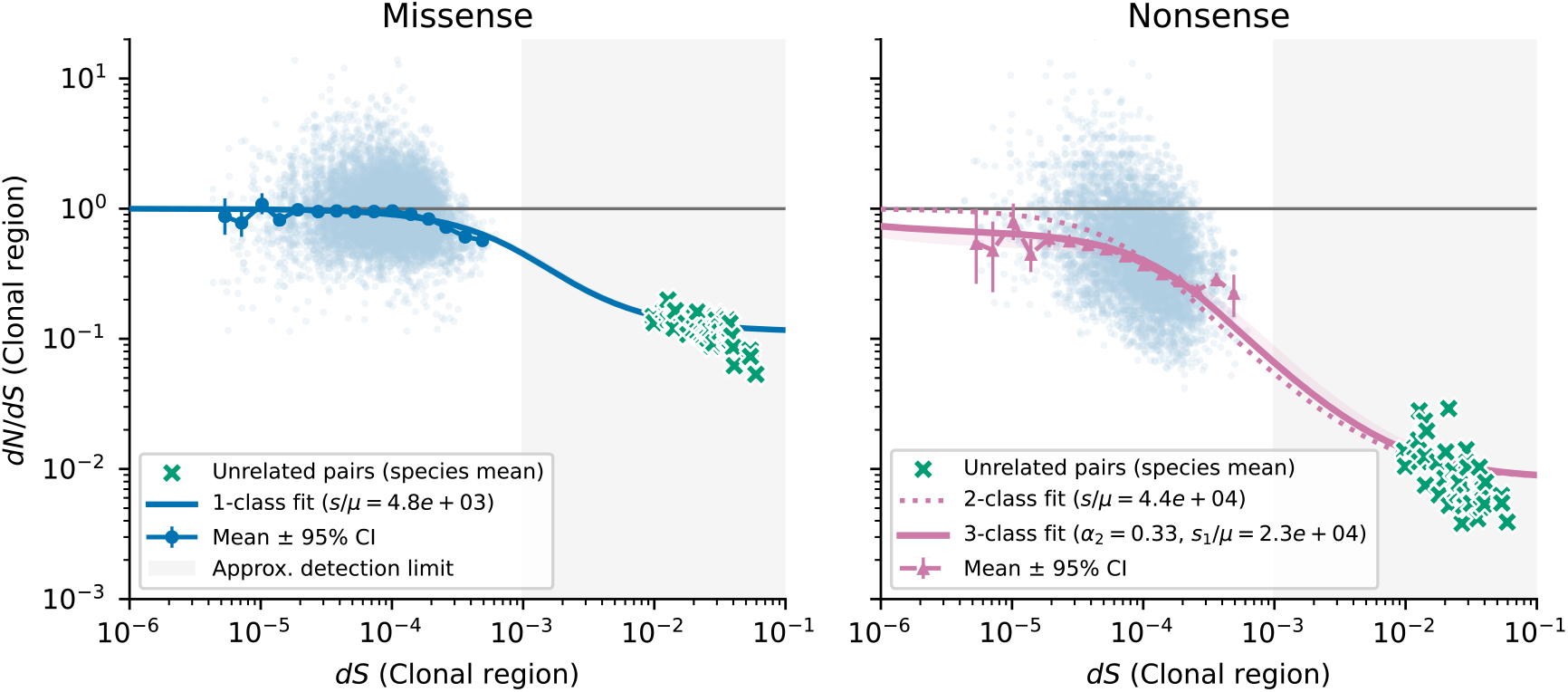
Analogous versions of Fig. 3 plotted for missense (left) and nonsense (right) mutations separately. In the left panel, the solid line denote the best-fit two-class purifying selection model (SI Section 4), with a 95% confidence interval smaller than the linewidth. In the right panel, the dashed line denotes the best-fit two-class model, while the solid line shows a three-class model with an additional class of strongly deleterious mutations with *s*_2_*/µ* ≈ 10^7^. These data suggest that nonsense mutations experience systematically higher fitness costs than missense mutations, but the average fitness cost of both mutation types remains comparatively weak.

In contrast, nonsense mutations showed a noticeably steeper decline in dN/dS with increasing divergence 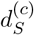, reaching a long-term value of ~ 0.01. This indicates that nonsense mutations are under stronger purifying selection, with a long-term neutral fraction of only ~ 0.01.

To quantify the underlying fitness costs, we fit these data to the same two- and three-class models that we used for the aggregate data above. The missense fits recovered nearly the same *s/µ* estimate as the aggregate data in Fig. 3. By contrast, we found that the nonsense mutations were better captured by the three-class model, which can account for *d*_*N*_ */d*_*S*_ *<* 1 at short times (Fig. 4; SI Section 4). The inferred parameters suggest that roughly 30% of nonsense sites fall into a strongly deleterious class with *s*_2_*/µ* ≫ 10^5^, which are effectively purged before the earliest observable divergence times in Fig. 4 (*d*_*S*_ ≪ 10^−5^). Given that the total target size for nonsense mutations is about 20-fold lower than missense mutations, these strongly deleterious variants correspond to only a few percent of all nonsynonymous sites — consistent with our earlier estimates from the aggregate *d*_*N*_ */d*_*S*_ dynamics in Fig. 3. Interestingly, for the remaining 60% of nonsense sites, the inferred selection coefficients are only about four-fold stronger than for missense sites, suggesting that many nonsense mutations are only weakly deleterious as well. These results demonstrate that stratifying dN/dS curves by mutation type can help resolve the heterogeneity in selection pressures across the genome: while most nonsynonymous mutations experience weak purifying selection, a subset of nonsense mutations are subject to strong constraint and are rapidly eliminated from the population.

### Comparisons to alternative models of adaptive reversion

While the purifying selection model captures much of the observed dN/dS dynamics within the clonal regions, it is not the only possible explanation. One intruiging alternative was recently proposed in Ref. 30, in which the nonsynonymous differences could instead arise from locally adaptive substitutions that revert on longer timescales when the local environment changes. Previous work has shown that this adaptive reversion model can produce clonal dN/dS trajectories with the same functional form as Eq. 1, despite arising from a different set of biological assumptions and parameter meanings (30; see SI Section 5 for a full derivation). To understand the implications of this alternative scenario, we sought to determine how the new recombination-corrected measurements in Figs. 3 and 4 further constrain the underlying parameters of the model.

For example, the speed of the decay in the adaptive reversion model is controlled by the reversion timescale *τ*_flip_, which is dominated by the time that it takes for the environment to shift to a state where the reverse mutation is preferred (30; SI Section 5). Using this mapping, the speed of the decay in Fig. 3 yields an associated reversion time of *µ* · *τ*_flip_ ≡ (*s/*2*µ*)^−1^ ≈ 4 × 10^−4^, which translates to *τ*_flip_ ~ 300 − 3000 years given current estimates of the mutation clock (*µ* ~ 10^−7^ − 10^−6^ per site per year; 8, 45). This revised timescale is significantly longer than a human lifetime, which places important constraints on the environmental processes that could be driving the shifts in the adaptive landscape.

A second key feature of the adaptive reversion model in Ref. 30 is that it predicts that the vast majority (~90%) of the clonal nonsynonymous SNVs in Fig. 3 must be comprised of reversible adaptive mutations. This large fraction of adaptive variants introduces an additional constraint linking the total number of adaptive sites and the degree of parallel evolution in other strains (SI Section 5). For example, if local adaptation was driven by a modest number of reversible adaptive sites, one would expect to see the same mutations independently arise in multiple pairs of strains. We quantified this signature of parallel evolution by identifying all purely clonal strain pairs (i.e. those with no detected recombination events) and asking how often the observed SNV differences between them were also found in other strains within our larger cohort (SI Section 6). Fig. 5A shows an example of this calculation for *Phocaeicola vulgatus*, one of the most prevalent gut commensals in our dataset. The data show that nearly 90% of all clonal nonsynonymous SNVs were private to a single clonal lineage and were never observed elsewhere among the 120 sampled hosts. Similar patterns are observed across the other species in our dataset (Fig. S8), with clonal SNVs rarely recurring in multiple strains (and at comparable rates between synonymous and nonsynonymous mutations). Stratifying SNVs by their recombination status is crucial for this calculation, since much higher rates of “parallelism” are observed when recombined sites are included (Figs. 5B and S8).

**Figure 5:**
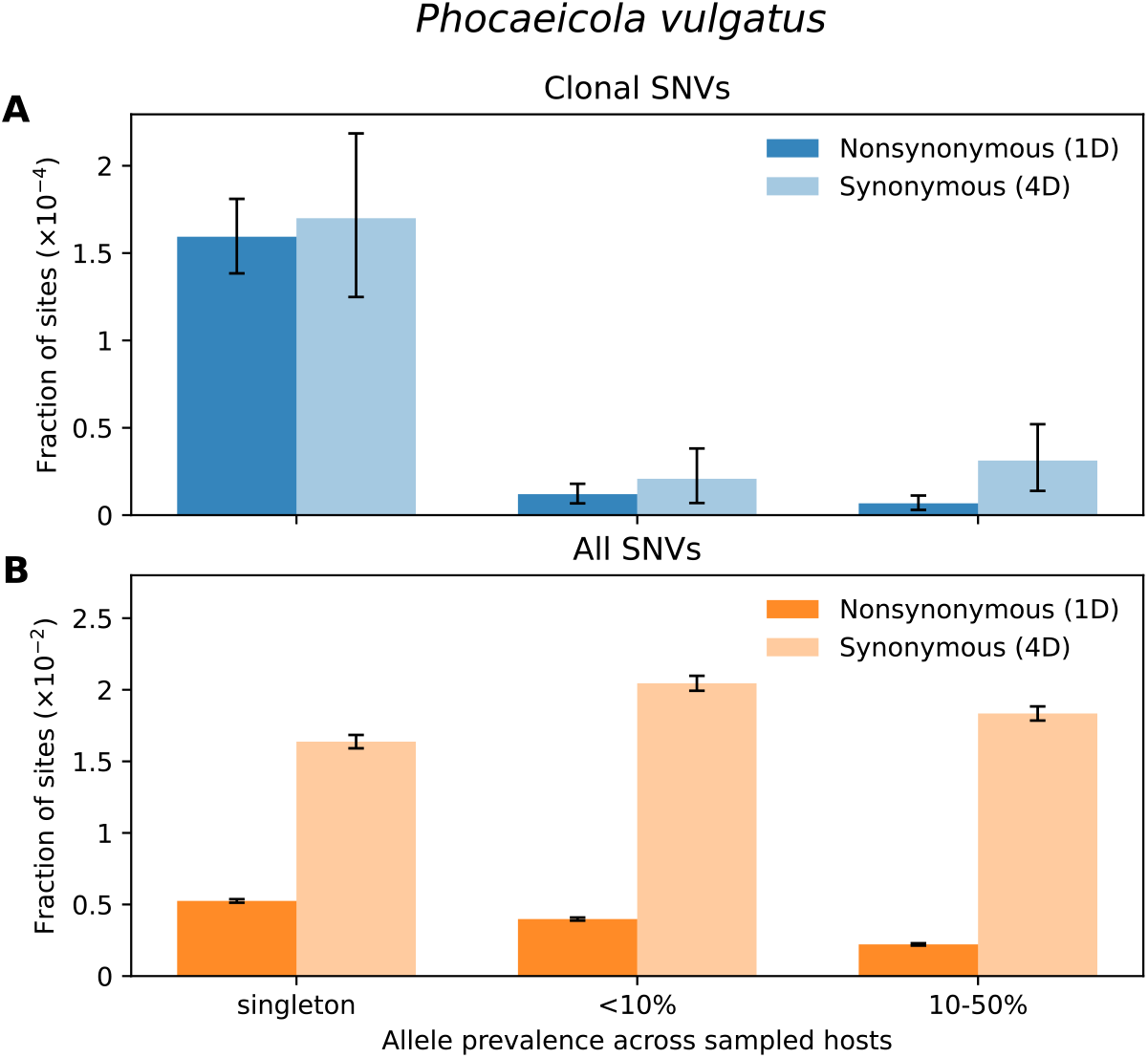
Measuring parallel evolution of clonal SNVs between closely related strains of *Phocaeicola vulgatus*. (A) Bar plots showing clonal SNVs in *P. vulgatus* stratified by the prevalence of the minor allele across other hosts in our cohort. Each bar shows the fraction of nondegenerate (1D; dark blue) or fourfold degenerate (4D; light blue) sites that harbor a SNV identified between fully clonal strain pairs (i.e., those with no detected recombination events). The singleton class refers to SNVs observed in only one of the 120 samples within the major clade of *P. vulgatus* (SI Section 6); other SNVs are grouped into two broad frequency ranges (<10% and >10%). These data show that most clonal SNVs are private to a single strain pair. (B) The same analysis applied to all SNVs in the dataset, most of which reside on recombined segments; the larger fraction of common alleles highlights the importance of stratifying SNVs by their recombination status. In both panels, error bars denote 95% confidence intervals assuming a Poisson distribution of observed SNV counts per bin. Analogous results for several additional species are shown in Fig. S8.

By extending the calculations in Ref. 30, we show that these low rates of parallel evolution can explain the dN/dS curves in Fig. 3 only if more than 1% of all nonsynonymous sites in the genome – corresponding to tens of thousands of sites – are targets of reversible adaptive evolution (SI Section 5). These numbers could be consistent with a smaller number of effective loci if mutations in the same locus interact with each other epistatically (e.g. loss-of-function mutations) (30). However, a similar calculation shows that at least ~500 adaptive loci, each containing tens to hundreds of sites, are required to match the observed dN/dS curves in Fig. 3 (SI Section 5). This new estimate is an order of magnitude larger than the ~50 adaptive loci previously estimated in Ref. 30, and reflects the ~10-fold longer reversion timescale implied by the recombination-corrected data.

Finally, while the purifying selection model mechanically predicts that *d*_*N*_ */d*_*S*_ → 1 at short times, the adaptive reversion model can only match this feature of the data if the fraction of adaptive sites and their corresponding substitution rates are precisely tuned so that their product is of order *µ* (SI Section 5). Moreover, this condition must separately hold within each species in our dataset, as well as for different classes of sites (e.g. the missense and nonsense mutations in Fig. 4), where both the decay rates and fractions of adaptive sites can differ. The mechanisms that could give rise to such fine-tuning currently remain unclear.

Interestingly, the parallelism analysis in Fig. 5 also reveals a small subset of clonal mutations that are observed at higher prevalence within the broader population. Some of these mutations likely correspond to shorter recombination events that were not detected by our HMM algorithm. However, we also note that the dN/dS ratios of these recurrent clonal SNVs are somewhat elevated relative to the population average (Fig. 5B), which captures the long-term expectation for recombined sites. Although the small number of prevalent clonal SNVs limits statistical power, this relative enrichment of nonsynonymous variants raises the possibility that a small minority of clonal mutations may still correspond to adaptive reversions, consistent with prior direct observations (28, 46, 47). Nevertheless, the small overall fraction of these recurrent clonal SNVs in Figs. 5 and S8 suggests they cannot be driving the large-scale temporal trends in Fig. 3.

## Discussion

The temporal dynamics of dN/dS carry important information about the selection pressures operating in natural bacterial populations. Here we have shown that in a broad range of human gut bacteria, the dN/dS ratios between closely related strains are dominated by the steady accumulation of recombination events, rather than the gradual unfolding of selective effects over time. By explicitly masking these recombination events, we demonstrated that the residual dN/dS levels in the clonally inherited portions of the genome remain informative, and can be used to constrain different models of selection. We found that a simple purifying selection null model can quantitatively reproduce the recombination-corrected dN/dS curves (Figs. 3 and 4), with estimated fitness costs that are about ~10-fold lower than previous analyses of the non-recombination-corrected data (28–30). The implied fitness costs in this model are remarkably weak (*s* ≪ 10^−4^), and would be difficult to observe on the timescales of a laboratory experiment or a typical human generation time. Nevertheless, these nonsynonymous variants are still reliably purged on the longer evolutionary timescales separating typical pairs of strains from different hosts (*s* ≳ 1*/T*_mrca_).

How natural selection is able to optimize such weak fitness costs is still a major open question — particularly given the pervasive hitchhiking and selective bottlenecks that are observed in the gut microbiome on shorter evolutionary timescales (8, 28, 48–52). However, it is worth emphasizing that the selection coefficients in Eq. 1 are really effective fitness costs (53), which average over multiple rounds of host colonization, growth, and dispersal. In metapopulation contexts like the microbiome, it is currently unclear how much of this selection is driven by competition between individual cells, competition between co-colonizing lineages, or larger-scale ecological processes (e.g. multi-level selection between hosts; 54). Understanding how the long-term dynamics in Eq. 1 emerge from these local evolutionary processes is an important direction for future research — one that will likely require new advances in modeling selection and genetic linkage in metapopulation contexts.

Our recombination-corrected data also shed new light on the proposal that the declining dN/dS levels could be driven by a small number of adaptive loci that undergo frequent local fixations and reversions. Although this adaptive reversion model can be modified to accommodate our new data, it requires a much larger number of adaptive loci (>500) as well as the fine-tuning of multiple small parameters to match the nearly neutral dN/dS levels observed at short times. These arguments suggest that adaptive reversions may not fully explain the global dN/dS patterns in Fig. 3. However, their impact on microbiome evolution is still worth exploring. Our observation that some clonal SNVs are frequently observed in other hosts (Figs. 5 and S8) could hint at a smaller number of genuine reversion events. If so, these variants could serve as useful entry points for identifying the functional targets of adaptive reversions. It is also possible that recombination itself could be an important source of adaptive mutations (55) (and subsequent reversions), even if the recombined segments are not included in the clonal dN/dS curves in Fig. 3 (SI Section 5). Explicitly accounting for such processes remains an important avenue for future work.

More broadly, our findings highlight a fundamental limitation in using dN/dS dynamics to study selection in natural bacterial populations. Because recombination rapidly erases clonal segments in many species of bacteria, the timescale over which clonal SNVs can be observed is sharply constrained. This leaves the critical regime, where dN/dS transitions from nearly neutral levels to its long-term equilibrium, as effectively unobservable with our present approach. To probe selection across these intermediate timescales, alternative population genetic measures such as the site frequency spectrum (56) or the decay of linkage disequilibrium (57–59) may provide complementary insights. Understanding how these population genetic observables map on to real-time evolutionary dynamics from longitudinal studies may ultimately help clarify how natural selection operates in this complex, multi-scale ecosystem.

## Acknowledgments

We thank T. Lieberman and P. Torrillo for clarifications on the adaptive reversion model, and J. Ferrare, S. Walton, O. Ghosh, and J. McEnany for comments and feedback on the manuscript. This work was supported in part by a Stanford Bio-X Bowes Fellowship (to Z.L.), NIH NIGMS Grant No. R35GM146949 (to B.H.G.), NSF grant PHY-2309135, the Gordon and Betty Moore Foundation grant no. 2919.02, and the Chan Zuckerberg Initiative DAF grant to the Kavli Institute for Theoretical Physics (KITP). B.H.G. is a Chan Zuckerberg Biohub – San Francisco Investigator.

## Data and Code Availability

All code used in this study, along with instructions for accessing the relevant datasets, are available at Github (https://github.com/zhiru-liu/dNdS_dynamics).

## Author contributions

Conceptualization: Z.L. and B.H.G.; theory and methods development: Z.L. and B.H.G..; analysis: Z.L. and B.H.G.; writing: Z.L. and B.H.G.

## Supplementary Methods

### 1 Metagenomic analysis of human gut commensals

The dN/dS measurements in the main text were obtained from collection of shotgun metagenomic data that have been analyzed in several previous works (28–30, 34). This dataset comprises of 932 fecal metagenomes from 693 individuals from North America, Europe, and China. All samples were processed using the same reference-based pipeline described in Ref. 28. In brief, we used the MIDAS software package (60) to align the raw sequencing reads to a panel of reference genomes representing common gut bacterial species, and identified single nucleotide variants (SNVs) based on the raw read pileups in each sample. In this work, we restricted our attention to SNVs in core genes (those that were present in more than 90% of samples in which the species was reliably detected). We used the quasi-phasing approach in Ref. 28 to identify intra-host populations in which the genotype of the dominant lineage could be inferred with a high-degree of confidence. Applying this method to the entire dataset yielded a collection of 5,416 strain genotypes from 43 species. Filtered genotype and coverage matrices for these quasi-phased genomes are provided in the Supplementary Data.

To identify synonymous and nonsynonymous variants, we focused on the subset of SNVs that were biallelic across the entire collection of quasi-phased genomes, ensuring that each site had a single putative ancestral allele and a single derived allele. Under the assumption that the major allele is ancestral, we used the reference genome sequence to determine site degeneracy (e.g., non-degenerate or fourfold) and to classify each mutation as synonymous, missense, or nonsense. These annotations were then used to tally the number of observed differences and opportunities in each mutation class.

#### Note on species names

The analyses in this study were performed using version 1.2 of the MIDAS reference genome database (60). Subsequent taxonomic updates have divided the *Bacteroides* genus into *Bacteroides* and *Phocaeicola* (61), leading to changes in species names relative to the original MIDAS database. For consistency, we retain the MIDAS species names in supplementary figures, while adopting updated nomenclature in the main text where appropriate. The corresponding name translations are listed below for reference:

| MIDAS DB v1.2 | NCBI Taxonomy Browser |
| --- | --- |
| *Bacteroides coprocola* | *Phocaeicola coprocola* |
| *Bacteroides massiliensis* | *Phocaeicola massiliensis* |
| *Bacteroides plebeius* | *Phocaeicola plebeius* |
| *Bacteroides vulgatus* | *Phocaeicola vulgatus* |
| *Eubacterium eligens* | *Lachnospira eligens* |

### 2 Estimating dN/dS ratios from pairwise comparisons

To estimate the synonymous and nonsynonymous substitution rates between a given pair of genomes, we used the observed sequence divergence at fourfold-degenerate (4D; synonymous) and non-degenerate (1D; nonsynonymous) sites. These divergences were estimated from the ratio of the total number of SNV differences (*k*_*i*_) and the total number of covered sites (*L*_*i*_) for each class of mutations. Since our goal is to examine how *d*_*N*_ */d*_*S*_ varies as a function of *d*_*S*_, it is important to avoid the mechanical correlation that arises when the sampling noise in *d*_*S*_ is shared between both the input and output variables. Following the Poisson thinning approach in Garud et al. (28), we randomly partitioned the synonymous sites into two disjoint sets of sizes 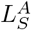 and 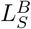, with corresponding difference counts 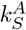 and 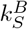. This yields a pair of synonymous divergence estimates for each a pair of strains,

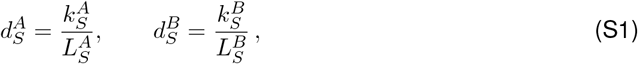

which should be statistically independent of each other after conditioning on the underlying divergence time. We therefore computed *d*_*N*_ */d*_*S*_ using 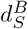 in the denominator,

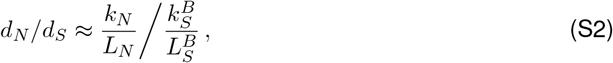

and plotted this ratio as a function of the independent variable 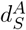.

When comparing closely related genome pairs from the gut commensal dataset, we further stratified sites by recombination status to compare *d*_*N*_ */d*_*S*_ between clonal and recombined regions. We used the recombination tracts that were previously inferred for these same pairs of strains in Ref. 34, using a pairwise hidden Markov model that leverages the spatial profile of synonymous sequence divergence across the genome. Closely related genome pairs were defined empirically as those sharing more than 50% identical genome blocks, where each block corresponds to 1,000 fourfold-degenerate sites. For each such pair, we used the inferred tract boundaries to assign each site to either the clonal or recombined class, and then computed *k*_*S*_, *L*_*S*_, *k*_*N*_, and *L*_*N*_ separately for each class before applying the Poisson thinning procedure described above.

While the above procedure provides *d*_*N*_ */d*_*S*_ for individual genome pairs, we also sought to compute the average *d*_*N*_ */d*_*S*_ value for all pairs of strains with a similar estimated divergence time. To do so, we first performed our Poisson thinning procedure on the clonal synonymous sites of each genome pair and used 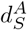 to assign pairs into logarithmically spaced divergence bins. Within each bin, we pooled all synonymous and nonsynonymous sites across pairs and computed a “ratio-of-means” estimate of the average *d*_*N*_ */d*_*S*_ within the bin:

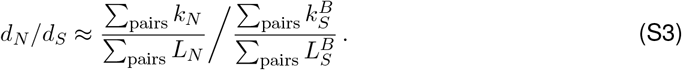

Confidence intervals were obtained from 600 bootstrap replicates, resampling both genome pairs and Poisson partitions. The same procedure was used to estimate the long-term *d*_*N*_ */d*_*S*_ for each species by analyzing a sample of 20 unrelated, fully recombined genome pairs.

Finally, to compare missense and nonsense substitution rates within clonal regions, we annotated the mutation class of each 1D SNV difference and calculated the total number of missense 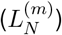 and nonsense 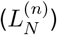 opportunities, where 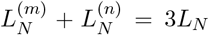. We then repeated the Poisson thinning and bootstrap procedures separately for each mutation class. In all cases, *d*_*N*_ */d*_*S*_ was calculated using the synonymous substitution rate per synonymous *opportunity*, while the *x*-axis in the relevant figures (e.g., Fig. 4) shows the synonymous substitution rate per synonymous *site*.

#### Note on transition/transversion composition

Partitioning 1D differences into missense and nonsense classes slightly changes the neutral per–opportunity mutation rate because the underlying transition:transversion (*Ti:Tv*) mix differs across these opportunity sets. In the standard genetic code, the Ti:Tv composition of missense opportunities is nearly identical to that of 4D sites, and therefore the expected neutral mutation rate per opportunity is effectively the same. Nonsense opportunities, however, are modestly enriched for transversions, with a Ti fraction of roughly 0.27 based on the genetic code. Assuming a typical bacterial rate bias of *µ*_Ti_*/µ*_Tv_ ≈ 4 (62), this composition would yield a neutral mutation rate per nonsense opportunity that is about 10% lower than at 4D sites. Thus, while this mechanism can produce a nonsense-only *d*_*N*_ */d*_*S*_ modestly less than one even under neutrality, values much lower than this expectation are indicative of the effect of selection.

### 3 Modeling the dynamics of genome-wide dN/dS with recombination

The phenomenological model in Fig. 2 assumes that each pair of bacterial genomes contains a mixture of clonally inherited and recently recombined genomic regions. We assume that in the clonal regions, sequence divergence arises solely through mutation, so the expected clonal synonymous divergence is given by

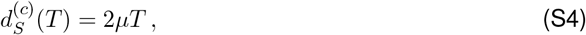

where *µ* is the per-site mutation rate and *T* is the divergence time between the two strains. In the simplest version of the model, we assume that the nonsynonymous clonal divergence is equal to 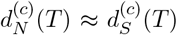, though we also consider more complex generalizations in Sections 4 and 5 below.

In contrast, the recombined regions of the alignment are assumed to originate from unrelated donors in the population. As a result, the sequence divergence within the recombined segments reflects the long-term evolutionary distance between unrelated strains. In the simplest version of the model, we assume that these recombined divergences are drawn from a single distribution with means 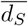 and 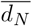, respectively, which can be directly measured from the larger dataset using the typical divergence between unrelated pairs of genomes.

The expected fraction of the genome that is covered by recombination events (*f*_*r*_) depends on divergence time *T* between each pair of strains. In our previous work (34), we showed that recombination dynamics in gut bacteria can be both heterogeneous across strains and nonlinear in time. To account for this complexity, we adopted an empirical approach to estimate the functional form of *f*_*r*_(*T*): we used our previously inferred values of *f*_*r*_ and 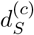 for each pair of closely related strains and fit these data to a logistic function of the form:

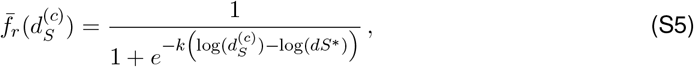

where *k* controls the steepness of the transition and *dS*^∗^ sets the characteristic divergence level at which recombination begins to dominate (Fig. S1). Using this functional form, the total expected divergence across the genome is a weighted average of clonal and recombined contributions:

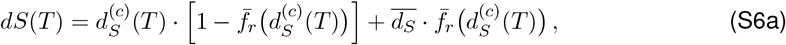

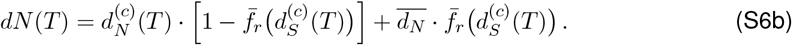

The ratio of these two expressions defines the expected genome-wide *d*_*N*_ */d*_*S*_ as a function of divergence time. This deterministic prediction is shown in Fig. 2B. In contrast to previous models (28, 30), which have free parameters (e.g. mutation rates, selection coefficients, and/or reversion rates) that must be estimated from the observed *d*_*N*_ */d*_*S*_ curve, the phenomenological model in Eq. S6 only depends on quantities that can be independently measured from other features of the data. Once 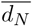, 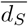, and 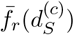 are specified, there are no additional free parameters for the theoretical curve in Fig. 2.

However, while this phenomenological recombination model provides an explanation for the observed decay of genome-wide *d*_*N*_ */d*_*S*_, it does not explicitly model the dynamics of *d*_*N*_ */d*_*S*_ within the clonal regions, or the values of 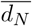 and 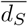 on the recombined segments. In the following two sections, we consider two classes of models that generate these predictions from different underlying mechanisms.

### 4 Purifying selection model

Multiple formulations of the purifying selection model describe how *d*_*N*_ */d*_*S*_ changes as a function of synonymous divergence (*d*_*S*_) at a locus evolving without recombination (22, 28, 30). These models converge on the common functional form in Eq. 1, which captures the impact of selection against nonsynonymous mutations with a distribution of deleterious fitness effects (DFE) *ρ*(*s*). The shape of this curve is governed by the ratio *s/µ*, which sets the timescale over which a mutation of effect *s* is purged. For an arbitrary distribution *ρ*(*s*), sites with different selection strengths are purged at different ranges of *d*_*S*_, and the *d*_*N*_ */d*_*S*_ curve reflects their relative contributions. Since the actual DFE is unknown, we used two different strategies to constrain this parameter from the observed data.

#### Estimating parameters for multi-class DFEs

Following previous work, we first considered discretized versions of the DFE, where nonsynonymous mutations fall into one of *K* discrete fitness classes with characteristic costs {*s*_*i*_} and relative target sizes {*α*_*i*_}. In the simplest version of this model with *K* = 2 classes, a fraction *α*_0_ of nonsynonymous sites are truly neutral (*s*_0_ = 0), and the remaining fraction *α*_1_ =1 − *α*_0_ share the same characteristic fitness cost *s*_1_. In this case, the *d*_*N*_ */d*_*S*_ curve reduces to

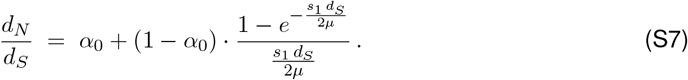

which depends on the two compound parameters *α*_0_ and *s*_1_*/µ*. At long times (*d*_*S*_ ≫ *µ/s*_1_), *d*_*N*_ */d*_*S*_ asymptotically approaches *α*_0_, which can be interpreted as the equilibrium fraction of neutral sites.

At short timescales (*d*_*S*_ ≪ *µ/s*_1_), the second term approaches 1 − *α*_0_, and thus *d*_*N*_ */d*_*S*_ → 1 regardless of *s*_1_, reflecting the fact that deleterious mutations have not yet been purged.

We also considered three-class versions of the model,

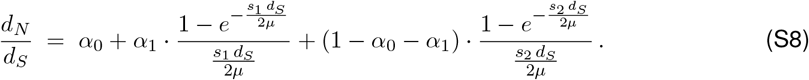

which include an additional fitness class with *s*_2_ ≥ *s*_1_. This three-class formulation can capture cases like Fig. 4B where the short-time *d*_*N*_ */d*_*S*_ is significantly below 1, indicating that a substantial fraction of nonsynonymous mutations are strongly deleterious.

We fit Eqns. S7 and S8 to the binned *d*_*N*_ */d*_*S*_ data using weighted nonlinear least squares. As described in SI Section 2, we used the Poisson–thinning procedure to partition the synonymous sites into two disjoint sets, “A” and “B”, with 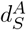 used for bin assignment and 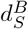 for the *d*_*N*_ */d*_*S*_ calculation. For each divergence bin *i*, let *K*_*N*,*i*_ and 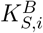 denote the total numbers of nonsynonymous and synonymous differences, respectively, obtained by pooling counts across all genome pairs in the bin for the “B” set of sites, and *L*_*N*,*i*_ and 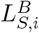 the corresponding pooled site counts. The binned *d*_*N*_ */d*_*S*_ estimate is then

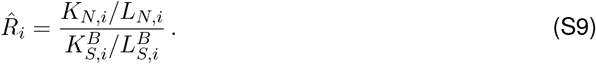

To account for varying statistical uncertainty across bins, we used inverse–variance weights,

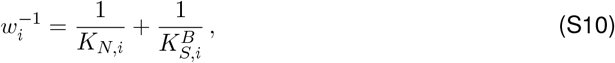

on a log-transformed version of 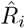 which down-weights bins with few observed differences. We excluded divergence bins containing fewer than three genome pairs to ensure that there was sufficient data for fitting. The resulting objective function is then defined by

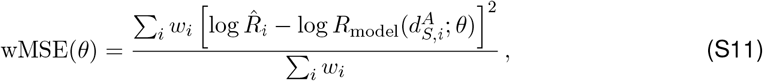

where 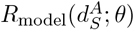 is the predicted *d*_*N*_ */d*_*S*_ from the chosen model (two-, or three-class) with parameters *θ*. For all model variants, we fixed the neutral-site fraction *α*_0_ to the long-term *d*_*N*_ */d*_*S*_ value for each species, estimated from unrelated genome pairs (SI Section 1).

We quantified the uncertainty in our estimates by bootstrap resampling the genome pairs, repeating the Poisson partitioning of the synonymous sites, and re-fitting the model to each bootstrap replicate. The 2.5%, 50%, and 97.5% quantiles of the bootstrap parameter distribution were used to define the corresponding confidence interval.

#### General constraints on the DFE from the asymptotic behavior of dN/dS

For an arbitrary DFE, the functional form in Eq. 1 still imposes strict constraints on the fraction of strongly deleterious mutations from the short-time behavior of *d*_*N*_ */d*_*S*_. A robust empirical observation across most of the species in our dataset is that the clonal *d*_*N*_ */d*_*S*_ curve remains close to one for 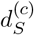 as large as 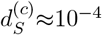. Intuitively, this implies that strongly deleterious sites cannot be too abundant: if many mutations were purged almost immediately, *d*_*N*_ */d*_*S*_ would already have dropped well below one at these short times.

We can formalize this argument by partitioning the integral in Eq. 1 at a user-specified threshold *s*^∗^, and then bounding the integrand in each term by its value at the lower limit of integration:

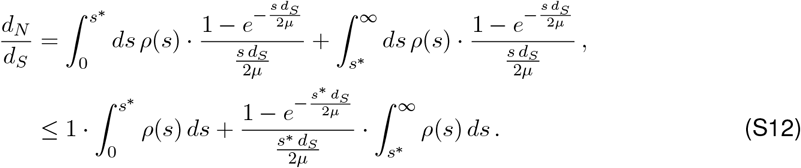

Rearranging this expression yields an upper bound on the fraction of mutations that are stronger than *s*^∗^:

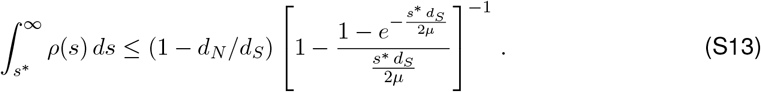

As a concrete example, Fig. 3 shows that *d*_*N*_ */d*_*S*_ ≈ 0.9 for *d*_*S*_ ~ 10^−4^. Eq. S13 then implies that more than 90% of nonsynonymous mutations must have *s/µ <* 10^5^ and more than half must have *s/µ <* 10^4^, independent of the detailed form of the DFE.

An analogous argument for the large *d*_*S*_ limit of the dN/dS curve yields a lower bound on the fraction of mutations that are more costly than a given *s*^∗^ value:

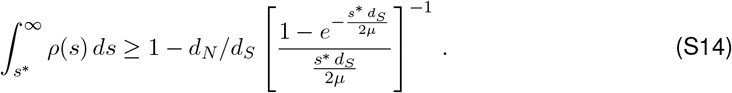

Applying this bound to the right-hand side of the *d*_*N*_ */d*_*S*_ curve in Fig. 3 shows that more than 80% of all nonsynonymous mutations must have *s/µ >* 10^2^, and more than half must have *s/µ >* 10^3^. Together, these two conditions provide general constraint linking the early- and late-time behavior of the *d*_*N*_ */d*_*S*_ curve to the underlying scale of the DFE.

### 5 Adaptive reversion model

We also compared the data in Figs. 3 and 4 to a generalized version of the adaptive reversion model in Ref. 30. This metapopulation model assumes that a subset of nonsynonymous sites undergo recurrent locally adaptive substitutions, which are later reversed when environmental conditions change.

#### Derivation of model

Following Ref. 30, we assume that these locally adaptive sites exhibit a form of modular epistasis. Each site is part of a larger module (or “locus” in Ref. 30) with two states that we denote as “mutant” (M) and “wildtype” (WT). We assume that mutations at any of the sites in the module will switch it from the wildtype to the mutant state, but compensatory mutations (M→WT) can only occur through a reversion of the original mutation. We also assume that there is strong intra-module epistasis, such that two or more mutations in the same module have the same phenotypic effect as a single mutation.

Substitutions and reversions within a single person’s microbiome are modeled as stochastic transitions with total rates *r*_on_ and *r*_off_, respectively. These two effective parameters encapsulate both the rate of environmental switching and the time that it takes for the population to reach the new locally adapted state. In many cases of interest, this latter timescale may be negligible, and *r*_on_ and *r*_off_ can be interpreted as switching rate between environmental states where either the mutant or wildtype version of the module is preferred. Ref. 30 focused on a symmetric version of the model where *r*_on_ ≈ *r*_off_ ≈ 1*/τ*_flip_; for completeness, we consider general values of *r*_on_ and *r*_off_ to allow for cases where the mutant state is preferred in only a small fraction of microbiomes (*r*_on_ ≪ *r*_off_).

Under these assumptions, the probability that a given module is in its mutant state at time *t* follows a simple Markov process,

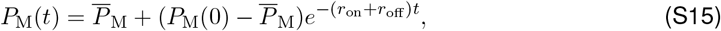

where 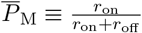 is the steady-state probability of finding the module in the mutant state. However, since dN/dS is a site-based metric, we must also understand the probability that a given *site* within the module will be found in its mutant or wildtype state. Under similar assumptions as above, we can model the dynamics at a given focal site as a three-state Markov process, corresponding to cases where (i) neither the module or the focal site are mutated (WT,WT); (ii) both the focal site and module are mutated (M,M); and (iii) cases where the module is mutated at another site within the module, while the focal site retains its wildtype allele (WT,M). The probabilities of finding the site in each of the three states evolve as

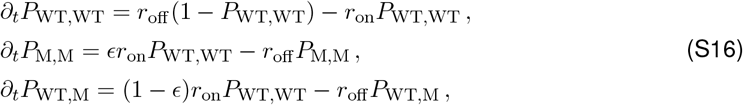

where *ϵ* is an additional parameter that describes the relative probability that a locally adapted module is driven by a mutation at the focal site. The solution to Eq. S16 is given by

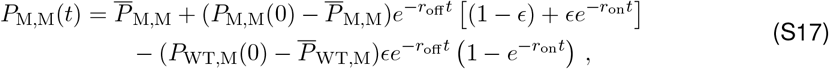

Where

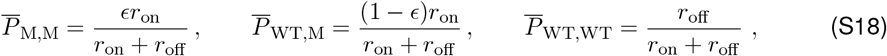

are the steady-state probabilities of finding the site in a given state. This expression reduces to Eq. S15 when each site comprises its own module (*ϵ* = 1), while allowing for more complex dynamics when *ϵ <* 1.

To calculate the expected genetic divergence between two independently evolving microbiomes, we assume that fixation within each microbiome is fast relative to the divergence time and that the shared ancestral allele is drawn from the steady-state multinomial distribution in Eq. S18. The expected probability of observing a difference at a given site is given by

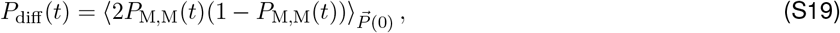

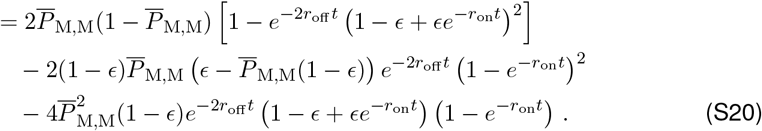

In our parameter regime of interest where each site makes a small contribution to its module (*ϵ* ≪ 1), this further reduces to

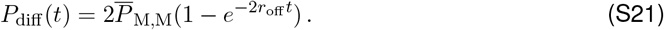

Summing over all nonsynonymous sites within a given focal set ℒ_focal_ yields a corresponding prediction for the *d*_*N*_ */d*_*S*_ ratio,

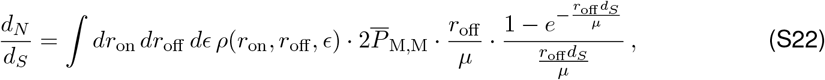

where *ρ*(*r*_on_, *r*_off_, *ϵ*) is the aggregate distribution of *r*_on_, *r*_off_, and *ϵ* across all of the sites within the focal set. This generalizes the derivation in Ref. 30 to allow for variable on and off rates, and applies to arbitrary subsets of nonsynonymous sites, in addition to the total genome-wide value. Notably, the functional form of Eq. S22 is identical to the purifying selection model in Eq. 1, even though the two expressions emerge from distinct evolutionary mechanisms.

#### Three-class model

To understand the constraints that this equivalence places on the underlying parameters of Eq. S22, we first considered a simplified version of the reversion model, similar to the one proposed in Ref. 30, where the nonsynonymous sites are divided into three discrete categories: (i) a set of effectively neutral sites with *r*_on_ ≈ *r*_off_ ≈ *µ*, (ii) a set of reversable adaptive loci with characteristic rates *r*_on_ and *r*_off_, and (iii) a set of strong, permanently deleterious sites with *r*_on_ ≈ 0 and *r*_off_ ≈ ∞. Denoting the fraction of sites in each class by *α*_0_, *α*_*r*_, and *α*_∞_ = 1 − *α*_0_ − *α*_*r*_, respectively, the *d*_*N*_ */d*_*S*_ curve in Eq. S22 reduces to

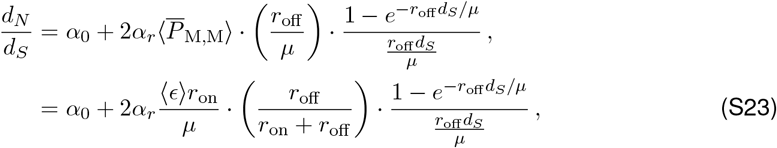

Where

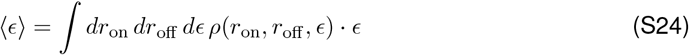

is the average value of *ϵ* across all of the sites within class (ii).

To more directly compare this expression with the one provided in Ref. 30, it is necessary to invoke some additional assumptions about the functional form of *ϵ*. We will assume that the probability that a given site will be the driver mutation of its module is proportional to its relative target size — a term we use in a generalized sense to refer to both its underlying mutation rate and any differences in its within-host fixation probability. Using this notation, we can express the value of *ϵ* for a given site *i* as

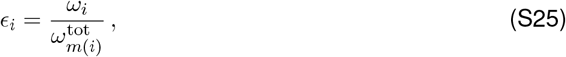

where *ω*_*i*_ denotes the relative target size for mutations at site *i, m*(*i*) denotes the module that site *i* belongs to, and 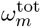 denotes the total target size for adaptive mutations in module *m*. This last term includes the contributions from (i) other nonsynonymous sites in the focal set, (ii) any nonsynonymous sites outside the focal set (if ℒ_focal_ is a subset of the genome), and (iii) any noncoding or non-point mutations (e.g. invertible promoters; 63) that can contribute to the overall adaptation of the module but are not included in dN/dS.

Summing over all of the nonsynonymous sites in class (ii), the average value of ⟨ *ϵ* ⟩ can then be expressed as

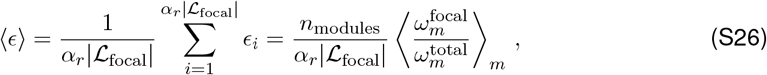

where *n*_modules_ is the total number of modules, 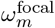 denotes the total target size of module *m* within the focal set, and the average in the angle brackets is now taken over all modules *m* that have at least one site in the focal set. Substituting this result into Eq. S23 yields a related expression for dN/dS:

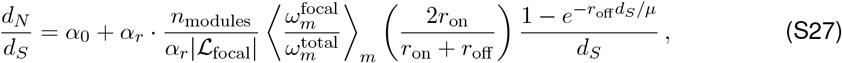

which reduces to the expression in Ref. 30 in the special case where *r*_on_ ≈ *r*_off_ ≈ 1*/τ*_flip_ and 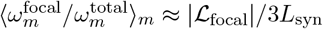.

In contrast to the purifying selection model in Eq. 1, all of the time-dependence in Eqns. S23 and S27 is generated by reversions of adaptive sites. This places strong constraints on the underlying parameters, which we can start to understand by considering the large and small *d*_*S*_ limits of this expression.

#### Comparisons to data

Like the purifying selection model in Eq. 1, the long-time behavior of Eqns. S23 and S27 is governed by the the fraction of effectively neutral nonsynonymous sites *α*_0_. In order to match the empirical observations of *d*_*N*_ */d*_*S*_ ≈ 0.1 between unrelated strains, we must continue to have *α*_0_ ≲ 0.1. Similarly, the timescale of the decay of *d*_*N*_ */d*_*S*_ suggests that the typical reversion rate must be on the order of *r*_off_ */µ* ~ 10^4^, analogous to the effective selection coefficient *s/µ* ~ 10^4^ inferred under the purifying selection model in Fig. 3.

One important difference between the two pictures is that while the purifying selection model was mechanistically constrained to approach one at short times, the adaptive reversion model possesses a different short-time limit,

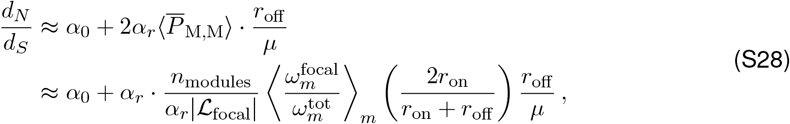

which functions as an additional fitting parameter. This extra degree of freedom allows the adaptive reversion model to produce dN/dS values that are substantially greater than one, which can be useful in cases where adaptive evolution is observed (8). However, for empirical curves like Figs. 3 and 4, where dN/dS remains comparatively close to one at short times, this additional degree of freedom must be precisely tuned in order to match this feature of the data. In particular, given the constraint that *α*_0_ ≲ 0.1, matching the empirical values of *d*_*N*_ */d*_*S*_ ~ *O*(1) at short times requires that the reversible component of Eq. S28 must also be order one, yielding a corresponding condition

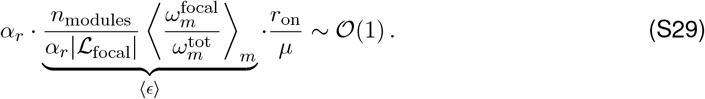

The left-hand side of Eq. S29 is a product of several independent parameters (*α*_*r*_, ⟨ *ϵ* ⟩, *r*_on_, and *µ*) which can differ from each other by several orders of magnitude in real biological systems. Achieving a dN/dS value of order one requires that these different terms must precisely balance one another. Since there is not an *a priori* reason why these different evolutionary parameters should be related to each other in this way, we argue that the adaptive reversion model requires an additional degree of fine-tuning to match this feature of the data.

Notably, this fine-tuning constraint must hold for any collection of sites within the genome that exhibit *d*_*N*_ */d*_*S*_ ~ 𝒪(1) at short times. For example, Fig. 4 shows that Eq. S29 must separately hold within the subsets of missense and nonsense mutations, even though the underlying parameters could differ between them. Such agreement is not automatically guaranteed by the model and would require an additional degree of fine-tuning.

To illustrate this idea, we recall that the speed of the dN/dS decline in Fig. 4 was ~4x faster for nonsense versus missense mutations, which we interpreted as a 4-fold higher fitness cost of nonsense mutations under the purifying selection model. In the adaptive reversion picture, this same difference in the dN/dS curves implies that the environmental switching rates must be 4-fold faster for nonsense vs missense mutations, which would require in turn that the nonsense mutations must arise from a systematically different set of modules than their missense counterparts. However, in order for the condition in Eq. S29 to remain satisfied in this new setting, the larger value of *r*_off_ */µ* must be balanced by a 4-fold reduction in the remaining terms (e.g. the number of modules per focal site or the relative on-vs-off rates) to ensure that *d*_*N*_ */d*_*S*_ ~ 𝒪(1) at short times. At present, the mechanisms that would allow for such fine-tuning remain unclear.

Even stronger tests of the adaptive reversion model are possible if one can identify the subset of reversible adaptive loci. In this case, the remaining, non-adaptive portion of the genome should provide a direct readout of the effects of purifying selection, which is assumed to be very strong in the version of the model in Eqns. S23 and S27. While it is difficult to fully enumerate the set of adaptive loci, several previous studies have performed conceptually related analyses by leveraging recurrently mutated genes in collections of isolates sampled from individual hosts (8, 64).

In one study in *Bacteroides fragilis* (8) (one of the species included in Fig. 2), this subset of recurrently mutated genes exhibited a dN/dS ratio of 6.0, indicating strong positive selection (8). However, the remaining portion of the genome retained a dN/dS indistinguishable from one, similar to the genome-wide patterns in Fig. 3. A similar analysis in the pathogenic bacterium *Burkholderia dolosa* (64) observed a dN/dS ratio of 18 in the most recurrently mutated genes, and a residual dN/dS that was measurably less than one (dN/dS≈0.63), but still larger than the long-term value of *α*_0_~0.1 estimated from Ref. 65. In both species, these dN/dS values imply that either the vast majority of mutations in the non-parallel genes (1 − *α*_0_*/*(*d*_*N*_ */d*_*S*_) ≳ 80%) must also be locally adaptive — and subject to the same fine-tuning constraints above — or that they must be eliminated on longer timescales through a purifying-selection-like process similar to Eq. 1. By contrast, no studies (to our knowledge) have identified a suitably large collection of sites with a short-term dN/dS ≲ 0.1, as predicted by the strong form of the adaptive reversion model in Eq. S27.

Together, these analysis show how comparing dN/dS curves for different classes of sites can provide additional information about the underlying selective forces that operate within a population. Extending this approach to other subsets of the genome would be an interesting avenue for future work.

#### Constraints from parallel evolution

An additional set of parameter constraints can be obtained by considering signatures of parallel evolution — i.e. the probability that the same adaptive mutations are independently sampled in unrelated lineages. In the model defined above, the degree of parallel evolution depends on the equilibrium frequencies 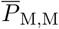 in Eq. S18, and how they vary across the different sites in our focal set.

To make this intuition more concrete, we recall that the short-time limit of Eq. S28 implies that 1 − *α*_0_*/*(*d*_*N*_ */d*_*S*_) ≈ 90% of all nonsynonymous substitutions at short times must be locally adaptive mutations. This suggests that any signatures of parallel evolution observed among these recent clonal mutations will provide additional information about the realized values of 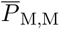. We implemented this idea using clonal strain pairs in *P. vulgatus* in Fig. 5 and found that the vast majority (*>* 90%) of nonsynonymous differences are private to a single strain pair (SI Section 6). For the adaptive reversion model to be consistent with this observation, the average steady-state probability 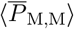 must be less than ~1*/n*, where *n* is the total size of the cohort. For our cohort size of *n* ~ 100, this implies that

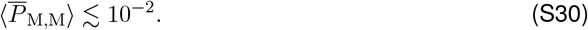

This constraint introduces a tension: if adaptive substitutions are rare because 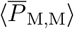 is small, then the fraction of adaptive reversible sites (*α*_*r*_) must be large to maintain the necessary nonsynonymous substitution rate. Combining this result with Eq. S28 and the estimated value of *r*_off_ */µ* ~ 10^4^ from Fig. 3, we find that *α*_*r*_ must satisfy the condition

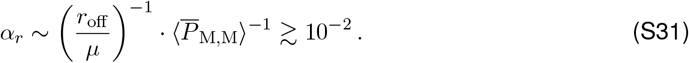

In other words, at least one percent of all nonsynonymous sites in the genome — corresponding to tens of thousands of sites — would need to participate in reversible adaptive dynamics to generate the observed *d*_*N*_ */d*_*S*_ pattern in Fig. 3. If recurrent clonal SNVs continue to remain as rare in cohorts of increasing sizes, the upper bound on 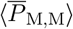 would become more stringent, requiring an even larger fraction of nonsynonymous sites to be reversible in order to explain the observed *d*_*N*_ */d*_*S*_ trend.

Similar arguments hold for more general distributions of *ρ*(*r*_on_, *r*_off_, *ϵ*). Echoing the purifying selection model in Eq. 1, the timescale of the *d*_*N*_ */d*_*S*_ curve requires that the vast majority of adaptive reversions have 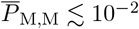 and 10^2^ · *µ* ≲ *r*_off_ ≲ 10^5^ · *µ*. These constraints suggest that the adaptive reversion model can account for the empirical *d*_*N*_ */d*_*S*_ patterns only if a substantial fraction of nonsynonymous sites experience reversible selection on environmental timescales much longer than the human generation time.

#### Limitations and future directions

Our analysis above has focused on a particularly simple form of the adaptive reversion model (Eq. S27), similar to the one employed in Ref. 30. Extensions of this model that incorporate more elaborate forms of epistasis, compensatory mutations (30), or mixtures of adaptation and purifying selection (59) could potentially some of the constraints identified above. Efforts to explore this space of models — similar to Ref. 30 — remain an important avenue for future work.

We have also neglected the effects of recombination. While this was motivated by our ability to focus on the clonal regions of the genome where the recombined segments have been removed, recombination can still affect this clonal diversity in an indirect manner by removing portions of the genome (and any variants they might contain) from both the numerator and denominator of dN/dS. If recombination events affect synonymous and nonsynonymous variants in an unbiased manner, their effects will cancel out when computing the clonal dN/dS ratio in Figs. 3 and 4. However, recombination could also be an important mechanism for generating the reversion events considered above (30), which could in principle lead to systematic biases in the realized recombination rates at synonymous versus nonsynonymous sites. In the simple models considered above, these biases would effectively contribute to the overall reversion rate *r*_off_. However, recombination can also be an important source of forward mutations, which would reduce the effective value of ⟨ *ϵ* ⟩ in the presence of modular epistasis, and further limit the contribution of reversible adaptive mutations to the clonal dN/dS curves in Figs. 3 and 4. Accounting for this behavior with more explicit recombination-aware models is another important avenue for future work.

### 6 Analysis of clonal versus population-wide SNVs

To examine the distribution of clonal SNVs across the broader host population, we focused on the subset of strain pairs that were predicted to be fully clonal (i.e. those with no inferred recombination events). Because multiple pairs of strains can all be closely related to each other, clonal SNVs can occur in more than one pair by common descent rather than by homoplasy. To avoid such redundancy, we filtered the initial strain set so that the remaining genomes were either clonal relatives or nearly fully recombined with respect to one another, defined as sharing less than 50% of their genome in identical blocks (SI Section 2). Within each group of strains connected by clonal relationships, we retained a single representative clonal pair. This procedure ensured that clonal variants were assessed only against otherwise unrelated genomic backgrounds.

In the case of *P. vulgatus*, our metagenomic dataset contains two major clades that are sometimes classified as separate species (*P. vulgatus* and *P. dorei*; 66). We restricted our analysis to the major *P. vulgatus* clade. After filtering, this procedure yielded a total of 120 strains, including 17 independent pairs of clonal strains. We catalogued all SNV differences at non-degenerate (1D) and fourfold degenerate (4D) coding sites between clonal pairs and computed the frequency with which each such variant appeared across the full set of 120 strains. For comparison, we also tabulated the prevalence of all SNVs present in at least one of the 120 strains, irrespective of whether they were originally identified in clonal pairs.

We repeated this procedure for each species with at least five clonal pairs, applying the same strain filtering and allele prevalence analysis. The resulting distributions of clonal versus population-wide SNVs, stratified by degeneracy class and allele prevalence, are shown in Fig. S8.

### 7 Analysis of *Staphylococcus aureus*

To assess whether our results extend beyond human gut commensals, we also analyzed a collection of 110 publicly available *Staphylococcus aureus* genomes obtained from Ref. 42 and originally sequenced by Ref. 67. These assemblies had already been aligned to the reference genome of strain MRSA252 in Ref. 42, and we annotated site degeneracy using the same reference–based in SI Section 1. When estimating *d*_*N*_ */d*_*S*_ between genome pairs, we utilized all sites jointly covered between the two genomes, rather than restricting to a predefined core genome set.

Unlike most gut bacteria, the recent evolutionary history of *S. aureus* is dominated by clonal expansion, with relatively few recombination events (42). However, three large hybridization events have been documented previously, corresponding to the formation of sequence types (STs) 34, 239, and 582. These strains arose through the replacement of ~200–600 kb regions in ST30, ST8, and ST25 genetic backgrounds, respectively (42–44). We used these previously identified tracts to stratify sites into “clonal” versus “recombined” categories and then estimated *d*_*N*_ */d*_*S*_ separately within each class using the same Poisson–thinning procedures described above (SI Section 2).

Pairs of genomes that did not belong to these hybrid sequence types were included in the genomewide *d*_*N*_ */d*_*S*_ analysis, but were omitted from the stratified clonal versus recombined comparison. Despite the limited number of recombination events, the resulting dynamics again revealed distinct trajectories in *d*_*N*_ */d*_*S*_ between clonal and recombined regions, consistent with the predictions of our recombination model.

## Supplementary Figures

**Figure S1:**
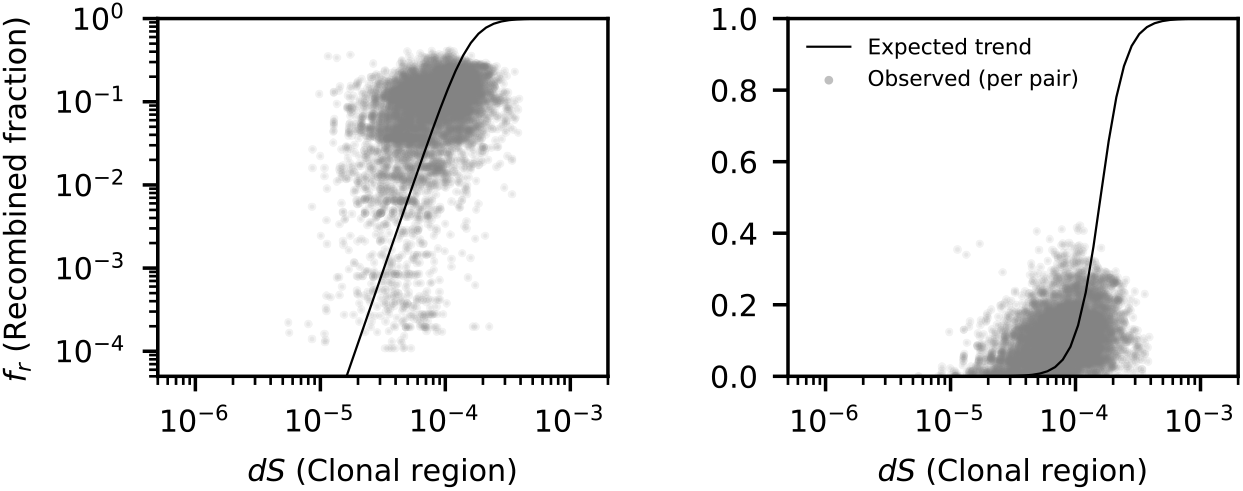
Empirical fit of recombination fraction versus clonal divergence. Each gray point shows the estimated recombination fraction (*f*_*r*_) and the associated clonal divergence (*dS*_*c*_) for each strain pair. The solid black line shows the logistic fit described in Eq. S5, with parameters *k* = 10 and *dS*^∗^ = 1.5 × 10^−4^ chosen by visual inspection of the data. This logistic form captures the nonlinear increase of recombination fraction with divergence, providing an empirical model for 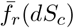 used in subsequent analyses. The two panels show the same data and fit, with the left axis on a logarithmic scale and the right axis on a linear scale.

**Figure S2:**
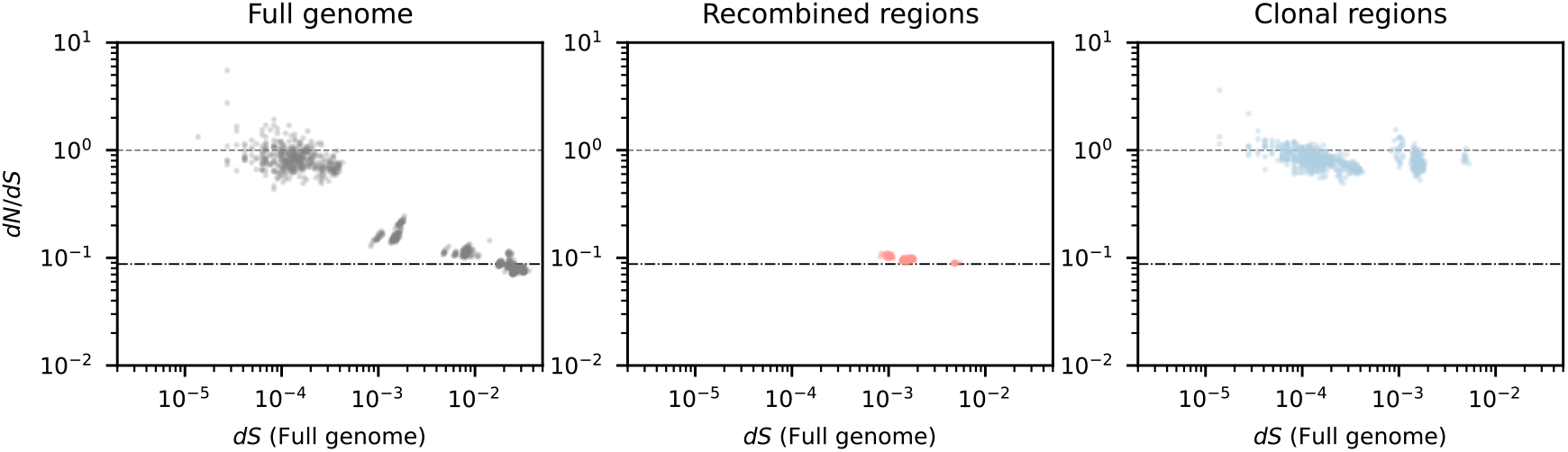
Analogous version of Fig. 2 for a collection of *Staphylococcus aureus* strains. Left: Genome-wide dN/dS dynamics across pairs of *S. aureus* strains, estimated from 1D and 4D sites. Middle: dN/dS values computed only within recombined regions corresponding to the large recombination events in ST34, ST239, and ST582 (SI Section 7). Right: dN/dS values computed within the remaining clonal regions.

**Figure S3:**
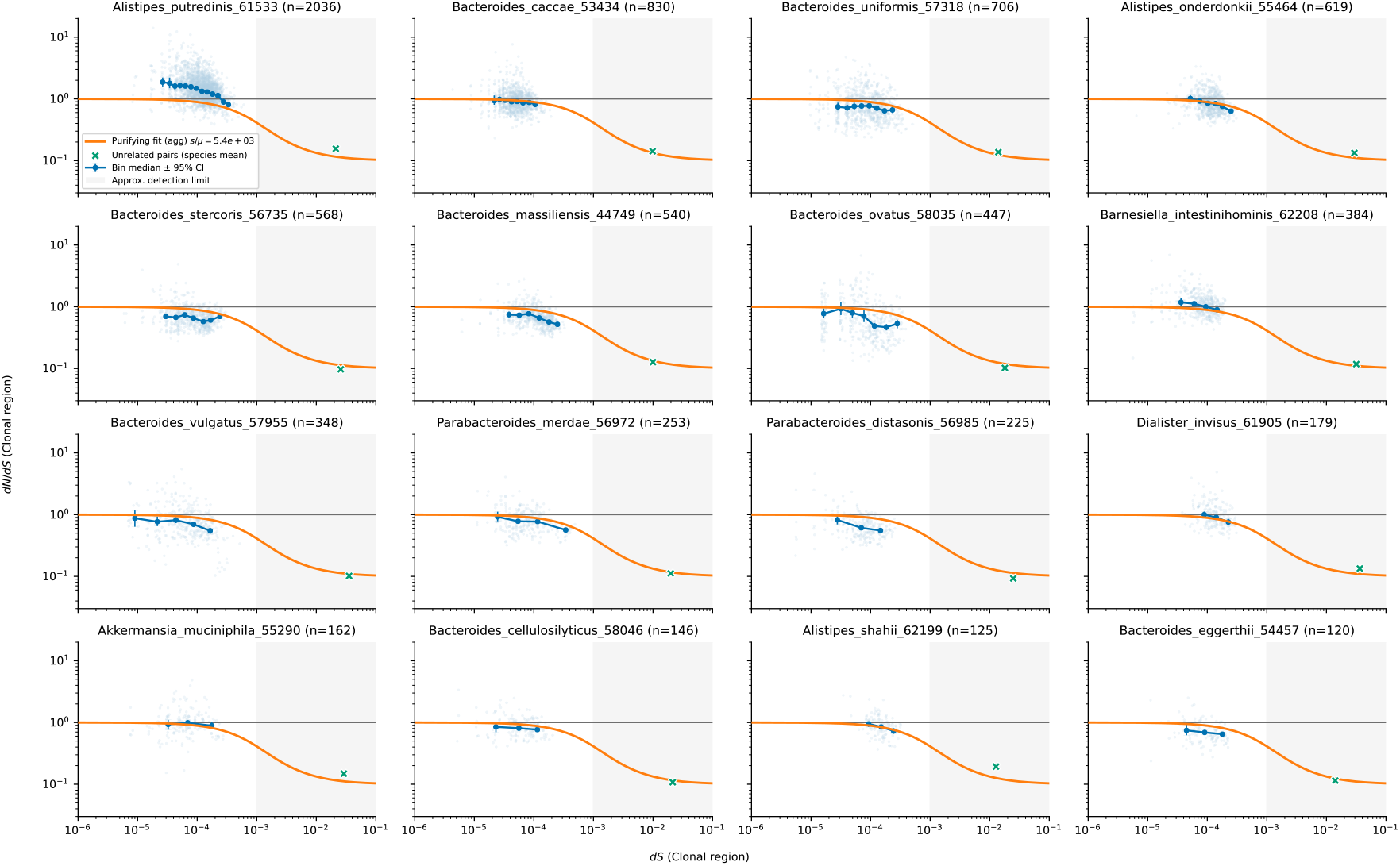
Species-level clonal *d*_*N*_ */d*_*S*_ dynamics (all nonsynonymous sites). Light-blue points show pairwise clonal *d*_*N*_ */d*_*S*_ versus clonal *d*_*S*_ estimated by Poisson-thinning; dark-blue markers with error bars denote the median of each divergence bin with 95% CIs from pooled counts. The orange curve is the same aggregate two-class purifying-selection fit used in Fig. 3, drawn identically in every panel for reference. Green “X” marks indicate the mean values of *d*_*N*_ */d*_*S*_ and *d*_*S*_ between unrelated, fully recombined pairs for that species; the gray line marks *d*_*N*_ */d*_*S*_ = 1; the shaded band denotes the approximate detection limit (*d*_*S*_ ≳ 10^−3^). Species and bins with too few pairs are omitted; panels are ordered by the number of available pairs in each species.

**Figure S4:**
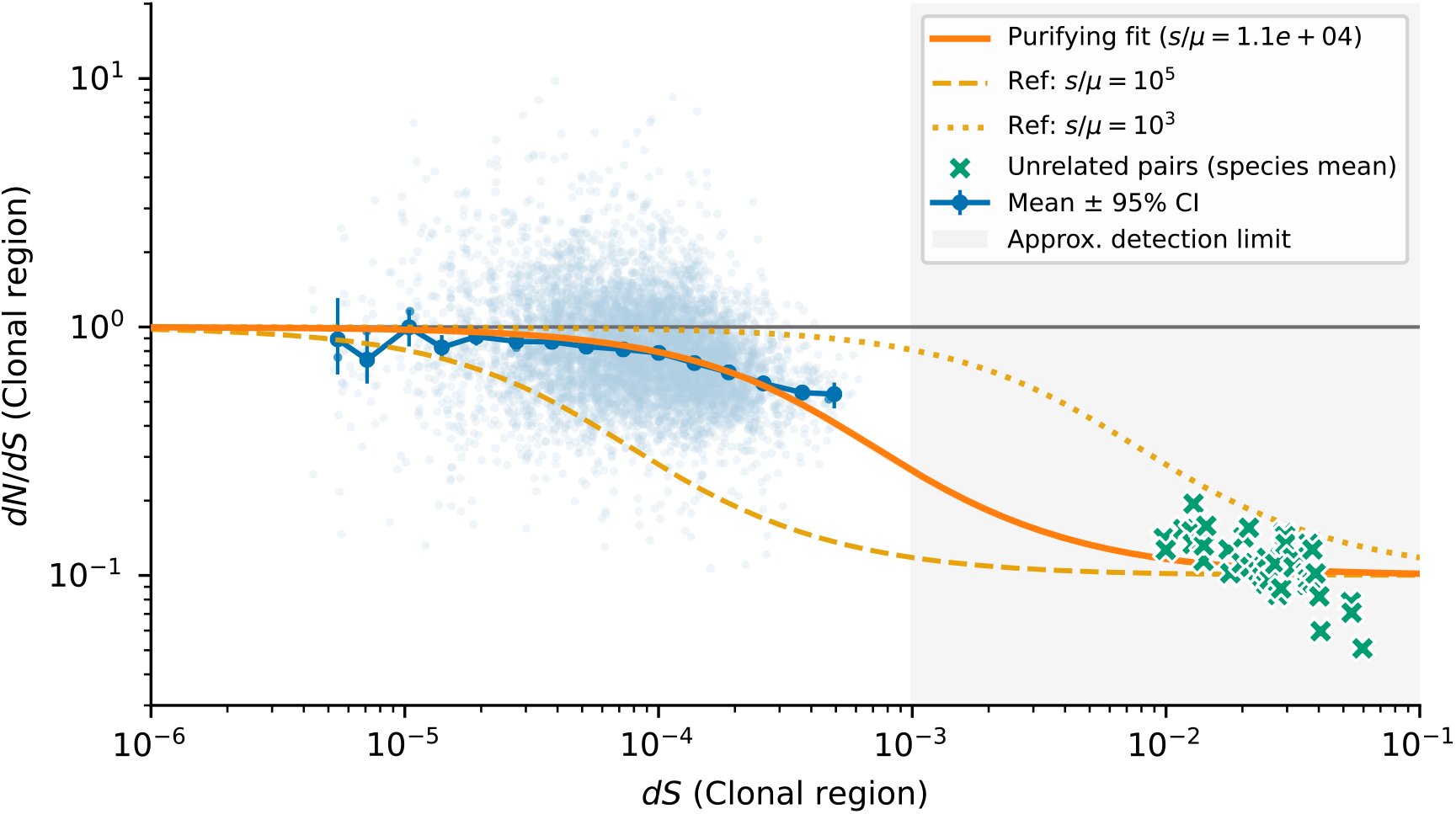
An analogous version of Fig. 3 excluding *Alistipes putredinis*. The overall results remain unchanged, with only a slightly higher estimated selection coefficient.

**Figure S5:**
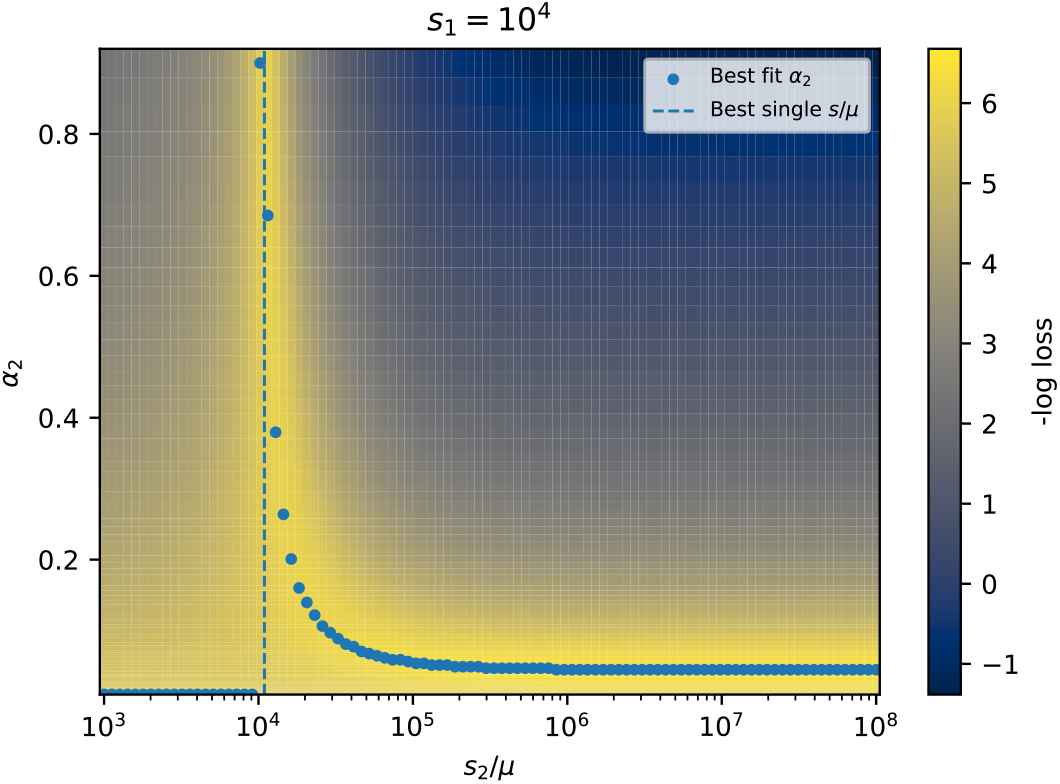
Loss surface of the three-class purifying selection model in SI Section 4. Each pixel shows the model fit quality (− log(wMSE), Eq. S11) as a function of the fraction of strongly deleterious sites (*α*_2_) and their scaled selection coefficient (*s*_2_*/µ*). Models were fit to all nonsynonymous sites, excluding *Alistipes putredinis*. The loss surface was evaluated by varying *α*_2_ and *s*_2_*/µ*, with *α*_0_ = 0.1 and *s*_1_*/µ* = 10^4^ held fixed at the values obtained from the best-fit two-class model. The dashed vertical line marks the best-fit single-class value of *s/µ*, while dots indicate the best-fit *α*_2_ value at each value of *s*_2_*/µ*. The shape of the loss surface suggests that introducing an additional strongly deleterious mutation class (*s*_2_*/µ* ≳ 10^5^) modestly improves the fit, but the inferred fraction of such sites remains very small (*α*_2_ *<* 0.05).

**Figure S6:**
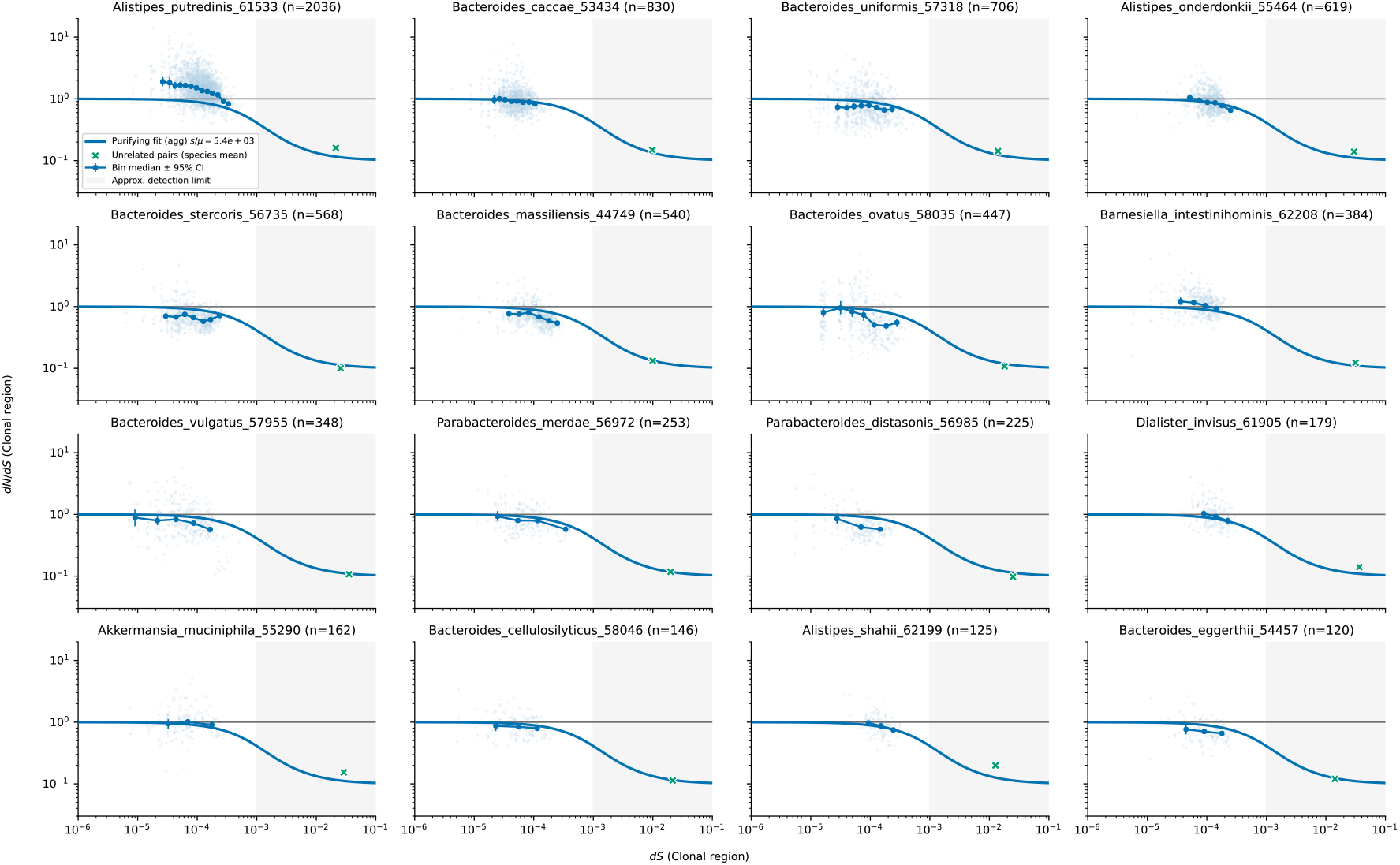
An analogous version of Fig. S3 computed for missense mutations only. The blue curve shows the aggregate two-class missense fit from Fig. 4 (left), held fixed across species.

**Figure S7:**
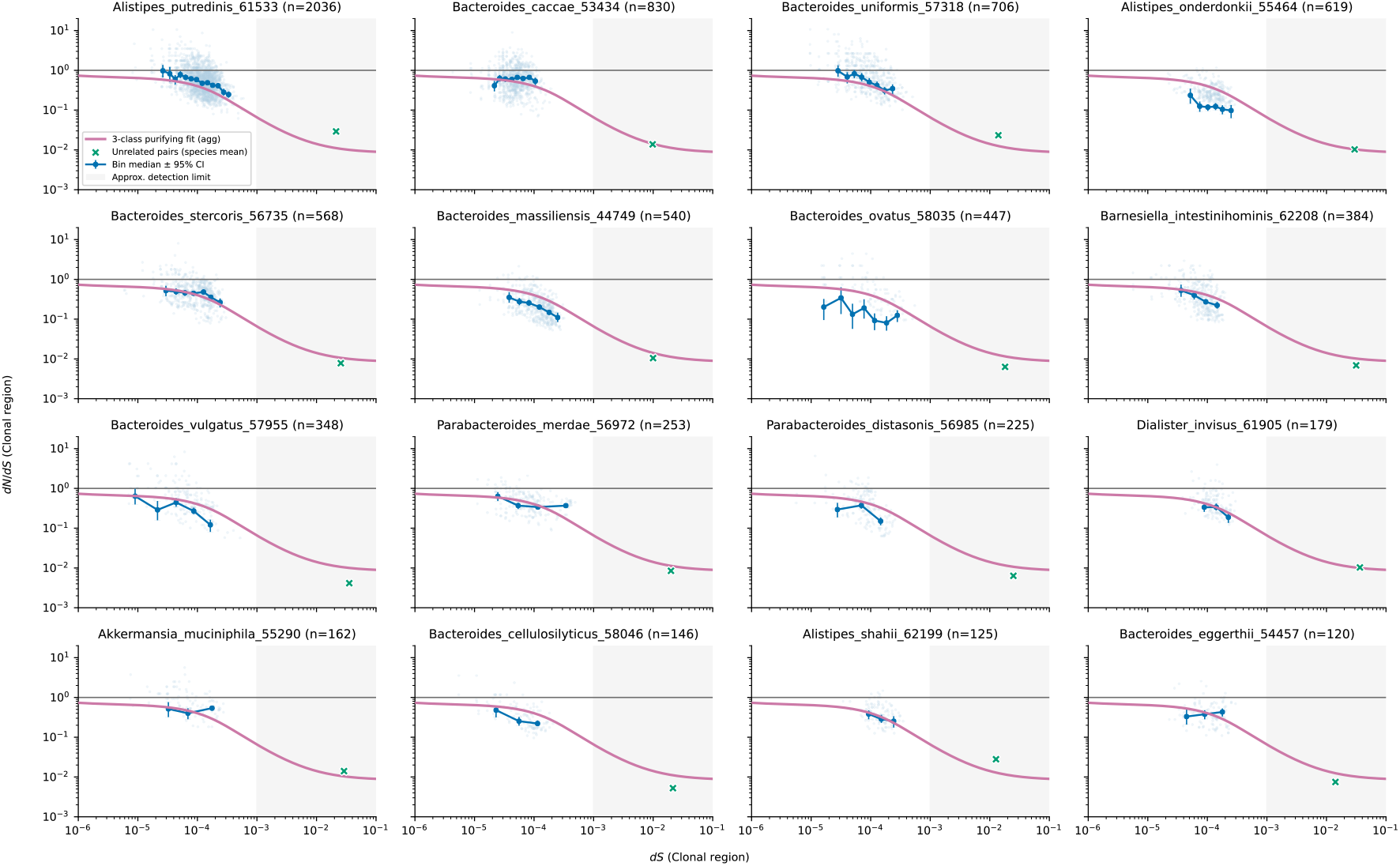
An analogous version of Fig. S3 computed for nonsense mutations only. The purple curve shows the aggregate three-class nonsense fit from Fig. 4 (right), held fixed across species.

**Figure S8:**
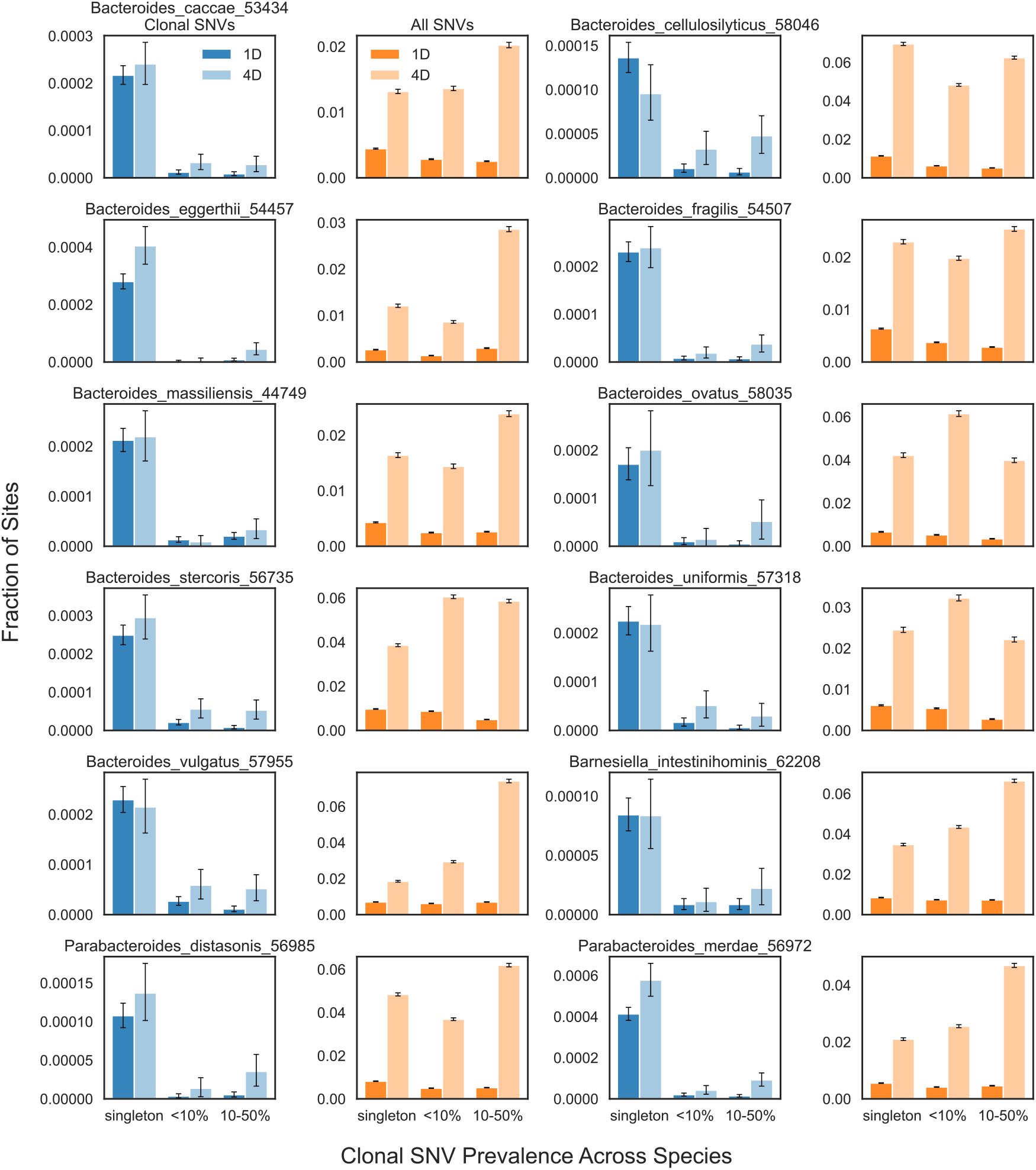
An analogous version of Fig. 5 for other commensal gut species. Left panels showing clonal SNVs (1D sites, dark blue; 4D sites, light blue) and right panels showing all SNVs for each species. “Singleton” denotes variants observed in only one host genome within each species. Error bars are 95% Poisson confidence intervals.

